# Acetylcholine disinhibits hippocampal circuits to enable rapid formation of overlapping memory ensembles

**DOI:** 10.1101/201699

**Authors:** Luke Y. Prince, Krasimira Tsaneva-Atanasova, Claudia Clopath, Jack R. Mellor

## Abstract

In the hippocampus, episodic memories are thought to be encoded by the formation of ensembles of synaptically coupled CA3 pyramidal cells driven by sparse but powerful mossy fiber inputs from dentate gyrus granule cells. Acetylcholine is proposed as the salient signal that determines which memories are encoded but its actions on mossy fiber transmission are largely unknown. Here, we show experimentally that cholinergic receptor activation suppresses feedforward inhibition and enhances excitatory-inhibitory ratio. In reconstructions of CA3 pyramidal cells, this disinhibition enables postsynaptic dendritic depolarization required for synaptic plasticity at CA3-CA3 recurrent synapses. We further show in a spiking neural network model of CA3 how a combination of disinhibited mossy fiber activity, enhanced cellular excitability and reduced recurrent synapse strength can drive rapid overlapping ensemble formation. Thus, we propose a coordinated set of mechanisms by which acetylcholine release enables the selective encoding of salient high-density episodic memories in the hippocampus.

## Introduction

The hippocampus plays a central role in the formation of episodic memories by processing information from the entorhinal cortex sequentially through the dentate gyrus, CA3 and CA1 regions. Anatomical, functional and theoretical considerations propose separate computational properties for each of these regions in support of memory processing (Marr, 1971; McClelland and Goddard, 1996; Kesner and Rolls, 2015). In particular, the CA3 region is characterized by a recurrently connected set of excitatory pyramidal neurons, which are believed to encode auto-associative memories by selectively strengthening recurrent synapses between ensembles of neurons that provide a neural representation of the memory (Nakazawa et al., 2002; Rebola et al., 2017). Configuring a recurrent network in this way is proposed to endow the network with attractor dynamics, in which the network is driven towards a stable state of ensemble formation (Marr, 1971; Hopfield, 1982; Kesner and Rolls, 2015). This process is related to memory retrieval, in which external sources of input will alter the state of the network by activating subsets of neurons and through recurrent dynamics will be driven towards these stable states in which all neurons in the ensemble are reactivated - a process also referred to as pattern completion (Gold and Kesner, 2005; Yassa and Stark, 2011). Within this framework, memory encoding is believed to be the procedure of altering the network through synaptic plasticity to create or change the position of attractor states (Treves and Rolls, 1994; Tsodyks, 1999). However, not all memories are stored, indicating that there may be a gate to select which experiences should be encoded, but it is unclear how such a filter might operate.

One potential filter mechanism is the release of neuromodulators such as acetylcholine that can rapidly reconfigure neuronal networks (Hasselmo, 2006; Prince et al., 2016). Acetylcholine is thought to promote encoding of new information by facilitating NMDA receptor function and induction of synaptic plasticity (Markram and Segal, 1992; Marino et al., 1998; Buchanan et al., 2010; Fernandez de Sevilla and Buno, 2010; Gu and Yakel, 2011; Dennis et al., 2016; Papouin et al., 2017) and selectively suppressing recurrent activity representing previously stored information in favour of feed-forward activity representing novel information (Hasselmo et al., 1995; Hummos et al., 2014). These properties are predicted to facilitate the encoding of new memories and allow greater overlap between representations (Hasselmo et al., 1995; Hasselmo, 2006). Commensurate with the concept of acetylcholine being important for encoding new information, acetylcholine release in the hippocampus and neocortex is associated with reward and arousal during periods of behavioural activity when it is beneficial to learn associated information in the environment to gain future rewards (Hangya et al., 2015; Teles-Grilo Ruivo et al., 2017). An alternative way to conceptualize the role of acetylcholine is as a signal for uncertainty, which may be resolved by familiarization through learning (Yu and Dayan, 2005). In either scenario, salience is indicated by increasing the release of acetylcholine in the hippocampus and other brain areas.

The dentate gyrus receives excitatory glutamatergic input from layer II of the medial entorhinal cortex, and sparsifies this signal by suppressing the activity of most dentate gyrus granule cells through lateral inhibition while dramatically increasing the firing rate of a select few granule cells (O’Reilly and McClelland, 1994; Leutgeb et al., 2007). By this mechanism granule cells detect salient, novel information and accentuate minor contextual details related to familiar information (a process often referred to as pattern separation). Individual granule cells provide a strong, sparse, facilitating input to a small number of CA3 pyramidal cells that can be sufficiently powerful to engage 1:1 spike transfer after multiple spikes in a granule cell burst (Acsady et al., 1998; Henze et al., 2002; Sachidhanandam et al., 2009; Vyleta et al., 2016). This focal excitation by mossy fibers drives synchronisation between subsets of CA3 pyramidal cells potentially allowing recurrent CA3-CA3 synapses to engage Hebbian plasticity mechanisms to create ensembles of strongly coupled CA3 cells thereby initiating the storage of new information (O’Reilly and McClelland, 1994; Treves and Rolls, 1994; Kobayashi and Poo, 2004; Brandalise and Gerber, 2014; Guzman et al., 2016; Mishra et al., 2016). Mossy fibers also excite a broad and diverse set of inhibitory interneurons that provide a widespread ‘blanket’ of feed-forward inhibition over a large population of CA3 pyramidal cells (Acsady et al., 1998; Toth et al., 2000; Mori et al., 2007; Szabadics and Soltesz, 2009). This feed-forward inhibition prevents runaway excitation and ensures tight spike timing for spike transfer (Torborg et al., 2010; Restivo et al., 2015; Zucca et al., 2017), while also enhancing memory precision (Ruediger et al., 2011) but it is not known what impact it may have on synaptic plasticity within the CA3 recurrent network.

Here we investigate the modulation by acetylcholine of ensemble creation and therefore memory encoding within the hippocampal CA3 network. Using slice electrophysiology and a hierarchy of experimentally constrained computational models of mossy fiber synaptic transmission and CA3 network activity, we demonstrate that acetylcholine suppresses feed-forward inhibition enabling plasticity at recurrent CA3-CA3 synapses and the formation of ensembles within the CA3 network. Furthermore, we show that acetylcholine increases the density of stable ensembles by enhancing permissible overlap between ensembles.

## Results

Acetylcholine regulates CA3-CA3 recurrent synaptic inputs and cellular excitability (Hasselmo et al., 1995; Vogt and Regehr, 2001; Dasari and Gulledge, 2011) which are both likely to be important for the creation of CA3 ensembles (Hasselmo et al., 1992). An additional critical feature of ensemble formation in CA3 is the mossy fiber input from dentate gyrus. However, the cumulative effects of acetylcholine on the mossy fiber projection incorporating both excitatory and inhibitory synaptic transmission are not known. Therefore, we first recorded experimentally the effect of acetylcholine on combined feed forward excitatory and inhibitory synaptic transmission in the mossy fiber pathway of mouse hippocampal slices. Minimal stimulation of granule cells resulted in EPSCs and IPSCs that were individually isolated by setting the membrane potential to -70mV and + 10mV respectively in accordance with experimentally determined reversal potentials for inhibitory and excitatory transmission respectively (Supplementary Fig. 1). Mossy fibers were stimulated with trains of 4 pulses at 20 Hz every 20 s. Application of the group II mGluR agonist DCG-IV (1 μM) reduced EPSC amplitudes by >90% (Fig. 1B) indicating selective activation of the mossy fiber pathway (Kamiya et al., 1996). The EPSC rise times (20-80%), latencies, and jitter were 0.57 ± 0.11 ms, 2.1 ± 0.49 ms and 0.53 ± 0.27 ms respectively (Supplementary Fig. 1) characteristic of mossy fiber synapses and confirming their monosynaptic origin (Nicoll and Schmitz, 2005). DCG-IV also reduced IPSC amplitudes by >90% and the rise times, latencies and jitter were 3.52 ± 1.08 ms, 6.20 ± 1.82 ms and 0.73 ± 0.25 ms respectively (Supplementary Fig. 1) indicating that IPSCs were mediated by disynaptic feed-forward inhibitory transmission in the mossy fiber pathway (Torborg et al., 2010). Both EPSCs and IPSCs exhibited pronounced facilitation in response to a train of 4 stimuli at 20 Hz as previously shown for mossy fiber feed forward excitatory and inhibitory pathways (Torborg et al., 2010).

**Figure 1:**
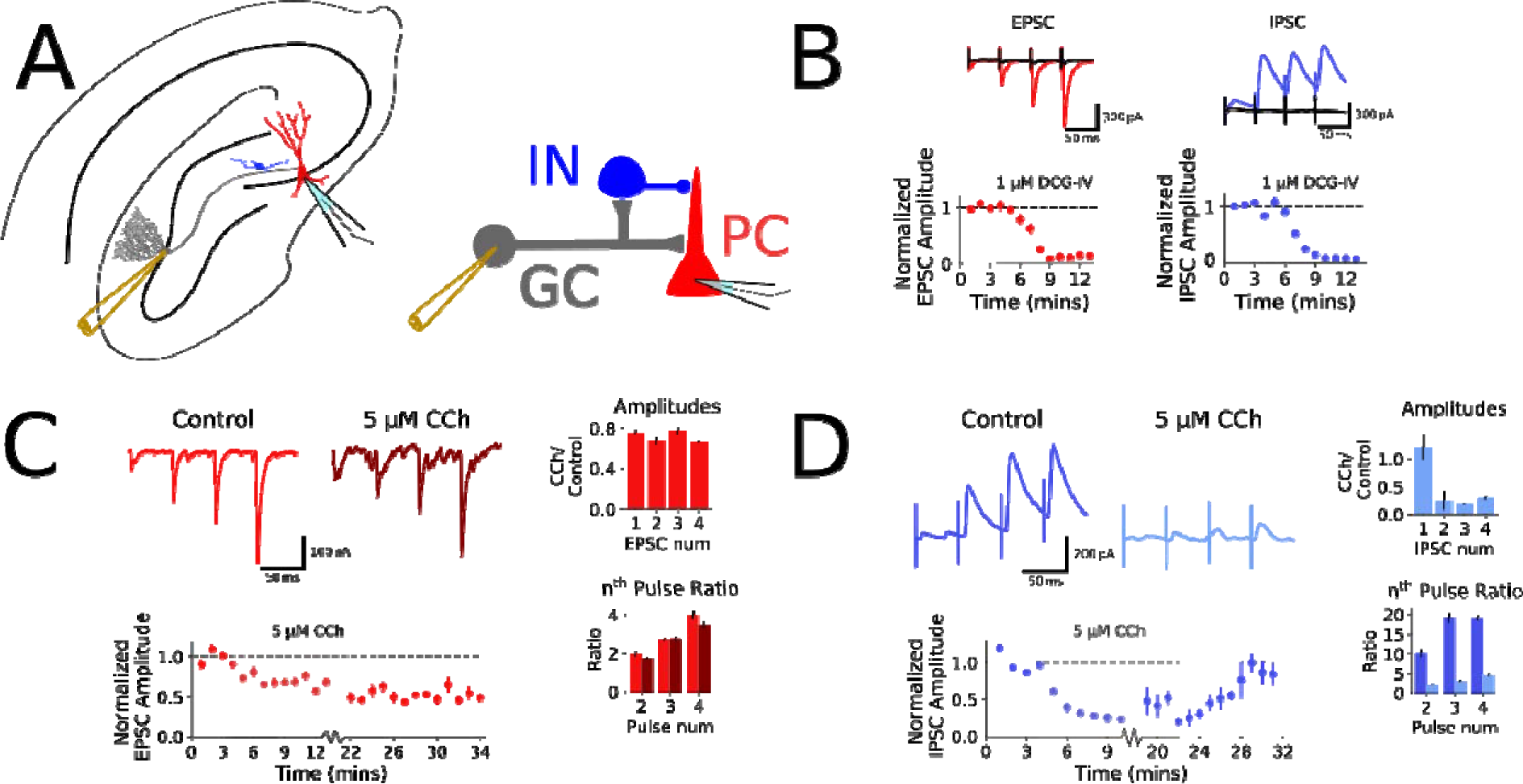
The effects of carbachol on feed-forward excitatory and inhibitory transmission in the mossy fiber pathway. A) Left: Experimental setup indicating the location of stimulation and recording electrodes within a hippocampal slice. Right: Schema of feed-forward mossy fiber circuit. B) EPSCs and IPSCs evoked by granule cells stimulation were blocked by 1 μM DCG-IV, confirming responses were driven by mossy fiber activation. Top: Example traces of DCG-IV block (black) of EPSCs (red) and IPSCs (blue). Bottom: Time course of DCG-IV block of 4th EPSC (red, n = 9) and IPSC (blue, n = 6). C) 5 μM CCh mildly suppresses mossy fiber EPSCs. Top left: Example traces before and after bath application of 5 μM CCh. Bottom Left: Time course of CCh effect, and washout (n = 7). Top Right: Effect of CCh on response amplitudes for each pulse. Bottom Right: Effect of CCh on nth/1st Pulse Ratio. D) 5 μM CCh substantially reduces disynaptic mossy fiber driven IPSC amplitudes. Top left: Example traces before and after bath application of 5 μM CCh. Bottom Left: Time course of CCh effect, and washout (n = 5). Top Right: Effect of CCh on response amplitudes for each pulse. Bottom Right: Effect of CCh on nth/1st Pulse Ratio.

### Acetylcholine reduces feed-forward inhibition in the mossy fiber pathway

The impact of acetylcholine on information transfer between the dentate gyrus and CA3 will depend on its effects on both excitatory and inhibitory pathways. To assess the effect of acetylcholine on both pathways we used the broad-spectrum cholinergic receptor agonist carbachol (CCh). Application of 5 μM CCh depressed EPSC amplitudes by ~25% (Fig. 1C; 75.6 ± 17.0% and 66.2 ± 11.9% of baseline measured at 1^st^ and 4th pulses, n = 7, *p* < 0.05) without altering facilitation ratios (Fig. 1C; *p* = 0.509; Measured at 1^st^ to 4^th^ pulse). This depression did not recover on washout of CCh. The use of minimal stimulation meant that responses to the first stimuli were highly variable and often very small or absent due to the low basal probability of release at mossy fiber synapses (Nicoll and Schmitz, 2005; Sachidhanandam et al., 2009; Torborg et al., 2010). This was particularly true for IPSC recordings resulting in very large facilitation ratios and a highly variable effect of CCh on the first IPSC in a train. In contrast to the effect on EPSCs, 5 μM CCh had no consistent effect on the 1^st^ IPSC in a train but depressed subsequent IPSC amplitudes reversibly and to a much greater degree (Fig. 1D; 121.5 ± 19.5% and 29.3 ± 13.5% of baseline measured at 1^st^ and 4^th^ pulses; n = 6, *p* = 0.012 measured at 4^th^ pulse) and at the same time reduced facilitation ratios (Fig 1D; *p* = 0.012; measured at 1^st^ to 4^th^ pulse). Because the larger IPSCs were greatly reduced by CCh this represents a substantial reduction in feed-forward inhibition across the 4 pulse stimulus train. CCh also enhances the excitability of neurons in the CA3 network (Hasselmo et al., 1995; Vogt and Regehr, 2001; Dasari and Gulledge, 2011) but this effect was absent at a cellular level in our recordings because of the inclusion of cesium in the pipette solution. Elevated network excitability was evident from a general increase in the frequency of spontaneous EPSCs and IPSCs (Supplementary Fig. 1). At lower concentrations, 1 μM CCh had limited effect on IPSC amplitudes whereas at higher concentrations 10 μM CCh had similar effects to 5 μM CCh with a substantial depression of IPSC amplitudes (Supplementary Fig. 1). These results indicate that acetylcholine causes a small irreversible depression of excitatory transmission at mossy fiber synapses whereas feed-forward inhibitory transmission is substantially depressed. Overall, this indicates a substantial net enhancement of Excitatory-Inhibitory ratio in the mossy fiber pathway in the presence of acetylcholine.

### Effects of acetylcholine on short-term plasticity at the mossy fiber synapse

Information transfer between the dentate gyrus and CA3 network depends on bursts of high frequency activity in dentate granule cells leading to pronounced frequency facilitation of excitatory synaptic input (Henze et al., 2002; Vyleta et al., 2016; Zucca et al., 2017). This is balanced by frequency-dependent facilitation of inhibitory synaptic input (Torborg et al., 2010) but variations in the short-term plasticity dynamics between the excitatory and inhibitory pathways will lead to windows within the frequency domain when excitation dominates and action potentials are triggered in CA3 pyramidal cells (Mori et al., 2004).

However, these temporal windows have not been fully characterized and, furthermore, the effect of acetylcholine on Excitatory-Inhibitory ratio over a range of mossy fiber stimulation patterns is not known. To investigate the patterns of activity that trigger action potentials under conditions of presence and absence of acetylcholine we adapted a Tsodyks-Markram based model of short-term plasticity dynamics in both excitatory and inhibitory pathways (see Methods).

Short-term plasticity models are difficult to constrain with responses evoked by regular stimulation protocols (Costa et al., 2013). Therefore, we constrained the model using responses to a stimulation pattern resembling the natural spike statistics of dentate gyrus granule cells which incorporate a broad range of inter-stimulus intervals (ISIs) (Fig. 2A) (Wiebe and Staubli, 2001; Mistry et al., 2011). Similar to the regular stimulation pattern of 4 stimuli at 20Hz, CCh depressed EPSCs and IPSCs in response to the irregular stimulation pattern across the range of ISIs but the depression was much more pronounced for IPSCs (Fig. 2B). Several short-term plasticity models of increasing level of complexity for both excitatory and inhibitory synaptic responses were assessed for fit to the experimental data (see Methods for detailed description of these models). The basic form of these models included a facilitation and a depression variable, here represented as *f* and *d* respectively. Dynamics for these variables are governed by parameters for degree (a) and timecourse of facilitation (*τ_f_*) and depression (*τ_d_*) as well as baseline of release (*f_0_*) and synaptic conductance (*g*) (Fig. 2B). Parameter inference for the short-term plasticity models was carried out and the best fitting models were selected by comparing the Akaike and Bayesian Information Criteria (AIC and BIC respectively) weights. These weights represent a normalisation of AIC and BIC values calculated by dividing the AIC and BIC values by the sum of these values across all models (log-likelihood of model given data punished for increasing complexity in two different ways). This is convenient as it allows these values to be transformed into a probability space and hence comparable across samples {Wagenmakers, 2004 #3194}. The model with the highest weight explains the data best. For excitatory mossy fiber synaptic transmission, a model containing a single facilitating variable with an exponent of 2 (*f^2^*) (Equations 1 and 2) best explained the experimental data (Fig. 2D).

**Figure 2:**
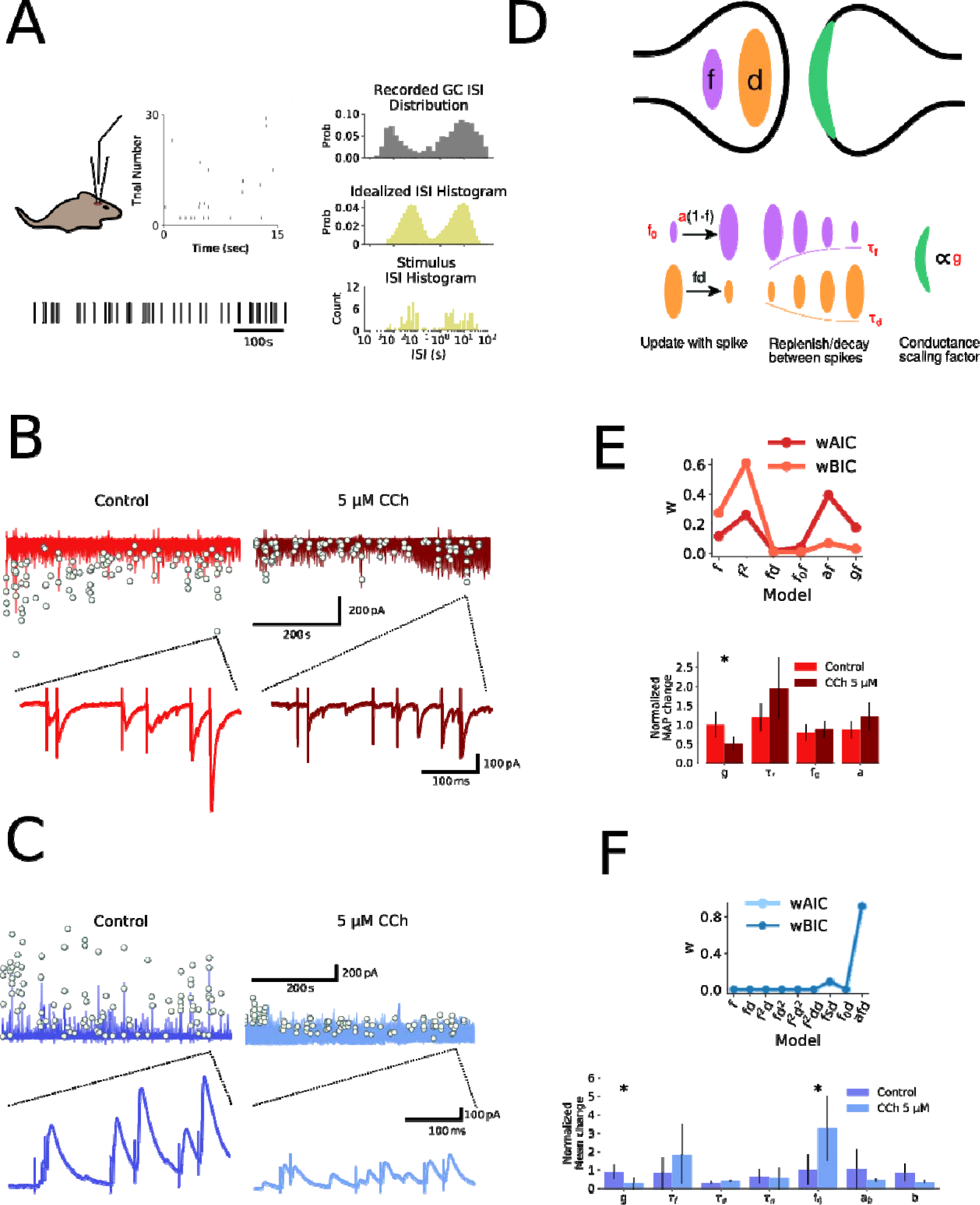
The mechanism of carbachol action on excitatory and inhibitory feed forward mossy fiber transmission determined by short-term plasticity models. A)Irregular stimulation protocol modelled on naturalistic granule cell spike patterns. GC spike patterns recorded *in vivo* during a spatial memory task, with a bimodal inter-spike interval (ISI) distribution (top right). Bimodal ISI distribution modelled as a doubly stochastic Cox process (middle right), with irregular stimulation protocol a sample drawn from this process (bottom right). B-C) Experimentally recorded EPSCs and IPSCs evoked by irregular stimulation protocol in hippocampal slices. Evoked peaks highlighted by white dots. An example burst is shown on expanded timescale. D) Tsodyks-Markram short-term plasticity model schematic illustrating facilitating (*f*) and depressing (*d*) presynaptic components with time constants (*τ_f_*, *τ_d_*) and postsynaptic scaling factor (*g*). E-F) Model selection and fitting for EPSCs (E) and IPSCs (F). Top: AIC and BIC weights for each fitted model. Model selection by highest AIC and BIC weights and evidence ratios. Bottom: Modulation of CCh assessed by effect on parameter fits normalized by time-matched control. * denotes significant parameter change.

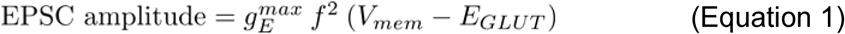

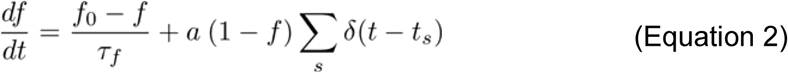

where *V_mem_* is the holding voltage of the cell in voltage clamp, 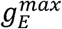is the maximum excitatory conductance, *E_GLUT_* is the reversal potential of glutamatergic transmission, and *t_s_* is the timing of the *s^th^* spike (or pulse). Explanation of other parameters for the short term plasticity model is given in the Methods section.

It is noticeable that AIC and BIC weights disagree on which model best explains the data. The *f ^2^* model had only the second highest AIC weight, but had the highest BIC weight, whereas the more complex *af* model had a higher AIC weight. However, the evidence ratio for BIC points favours the *f ^2^* model (P(*f ^2^*|Data)/P(*af* |Data) = 8.92 (see (Kass and Raftery, 1995)), whereas the evidence ratio for AIC indicates little evidence in favour of the *af* model (P(*af* |Data)/P(*f ^2^*|Data) = 1.51). Together this indicates that the *f ^2^* model best explains the data.

For inhibitory feed-forward mossy fiber synaptic transmission, a complex model with facilitation (*f*), depression (*d*), and additional facilitation over the increment parameter *a (afd*) produced the best fit (Fig. 2E and Equations 3, 4, 5 and 6) with both AIC and BIC weights convincingly pointing to the *afd* model as most appropriate to describe the data. Additional parameters in this model included a time course for facilitation of *a* (*τ_a_*), an increment scaling factor for *a* (*b*), and baseline (*a_0_).*

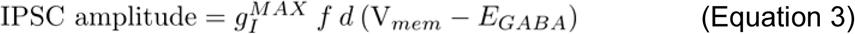

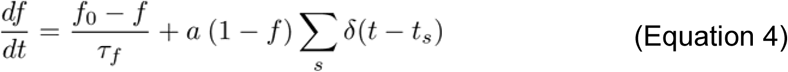

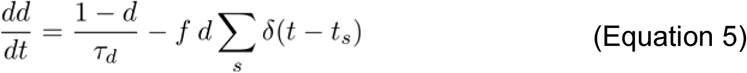

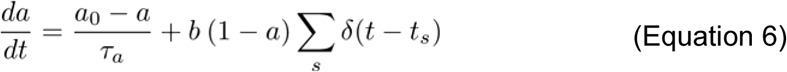

where 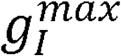 maximum inhibitory conductance and *E_GABA_* is the reversal potential of GABAergic transmission. See Methods for detailed explanation of the above equations.

Discrepancies between samples drawn from posterior-predictive distributions of these models indicated good fit for both models (Supplementary Fig. 2).

Using these models for the activity-dependent progression of excitatory and inhibitory synaptic weights we were then able to investigate the effect of acetylcholine on short-term plasticity by comparing normalized parameter estimates to time matched controls. Since posterior distributions for EPSC data were narrow and unimodal, maximum a posteriori (MAP) estimates were used, whereas mean parameter estimates were used for IPSC data since posterior distributions were wide and bimodal in some cases (Supplementary Fig. 2). This analysis revealed the small decrease in EPSC amplitude caused by CCh resulted from a reduction in the conductance scaling parameter ‘*g*’ (Fig. 2D; 49.9 ± 17.6%) in agreement with the data in Fig. 1C and indicating a postsynaptic mechanism. The substantial decrease in IPSC amplitude caused by CCh resulted from a large reduction in the conductance scaling parameter ‘*g*’, and an increase in the baseline parameter *‘f_0_’* which also had the effect of reducing facilitation (Fig. 2E; 73.1 ± 27.9% decrease in ‘*g*’; 225.7 ± 160.1 % increase in *‘p_0_’*). Since IPSCs are disynaptic, it is not straightforward to interpret how these parameter changes reflect biophysical changes to synaptic transmission, but the most likely explanation is a combination of increased feed-forward interneuron excitability and spike rate, coupled with a strong depression of GABA release. These results indicate the mechanisms underlying the effects of CCh on mossy fiber synaptic transmission and enable investigation of the granule cell spike patterns that favor excitation over inhibition.

### Enhancement of mossy fiber Excitatory-Inhibitory balance by acetylcholine

Feed-forward inhibition dominates excitation in the mossy fiber pathway for the majority of spike patterns (Mori et al., 2004; Torborg et al., 2010). Since acetylcholine depresses inhibitory transmission more than excitatory transmission (Figs 1 and 2) it is expected that the Excitatory-Inhibitory balance will be shifted towards excitation but the precise spike patterns that this occurs for are unclear. To examine how Excitatory-Inhibitory balance is affected by carbachol with different spike patterns we first tested the dependence of short term synaptic dynamics on background firing rate using the model for short-term plasticity dynamics. Spike patterns were described by two parameters: a between burst interval Δt*_between_* describing a background firing rate, and a within burst interval *Δt_within_* describing the time between spikes in a burst. The steady state value of *f, d,* and *a* given *Δt_between_* were then used to replace their baseline values (*a_0_ → a_∞_; f_0_ → f_∞_; d_0_ = 1 → d_∞_*) to set their initial values at the beginning of a burst, i.e.,

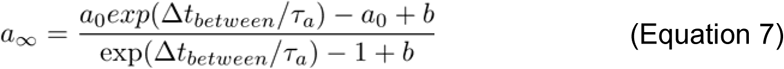

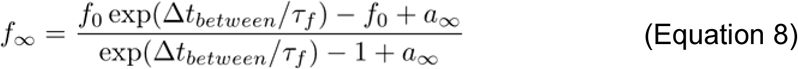

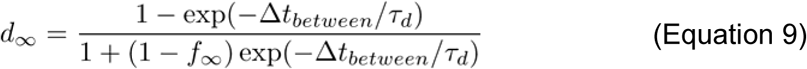

By systematically varying the between and within burst intervals for both excitatory and inhibitory synaptic input we were able to simulate EPSCs and IPSCs in the presence and absence of acetylcholine (Fig. 3A). The amplitudes of these responses were then used to explore the effects of acetylcholine on within burst facilitation ratios and Excitatory-Inhibitory balance across a wide range of between and within burst intervals.

**Figure 3:**
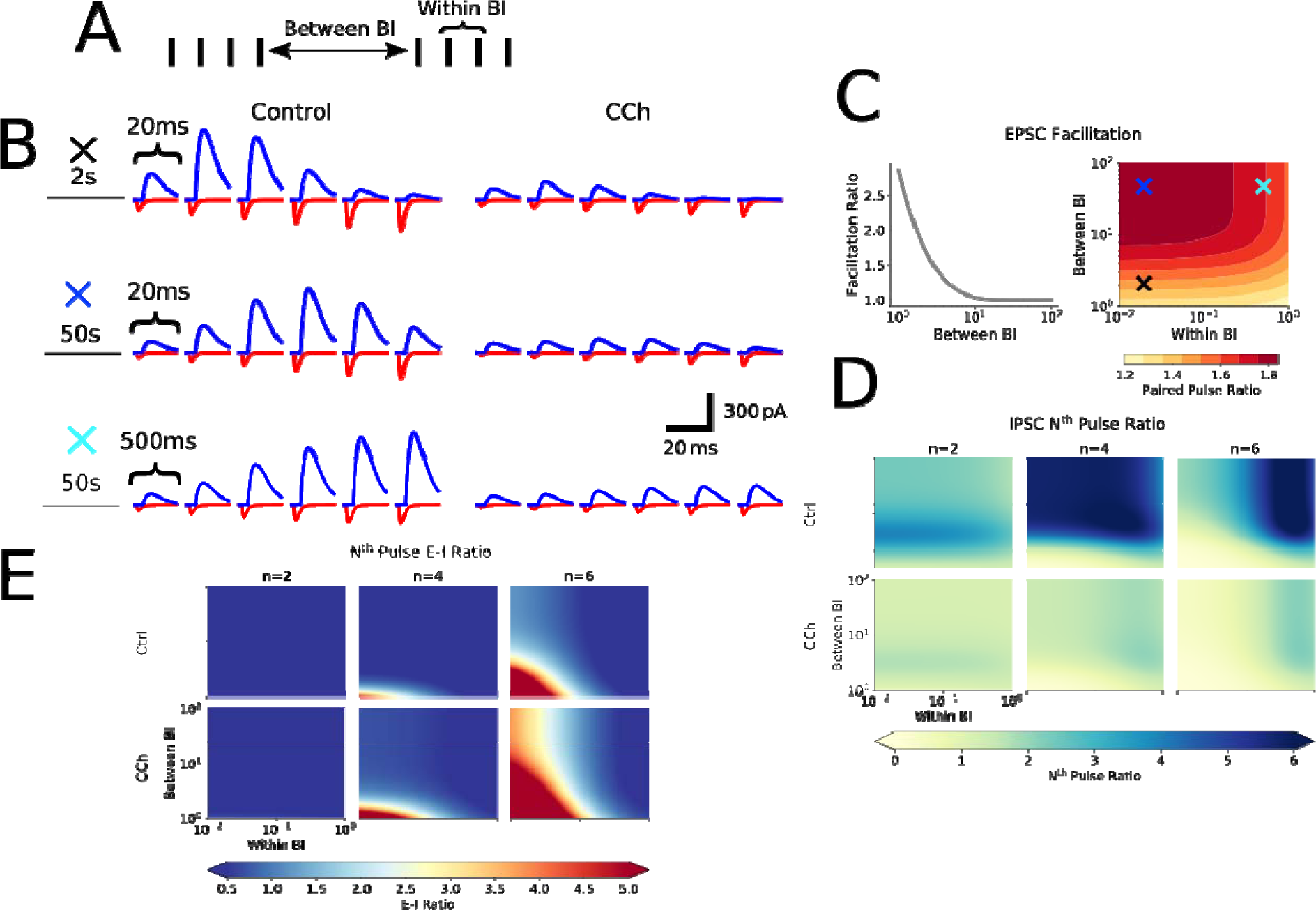
Carbachol alters the Excitatory-Inhibitory ratio within the feed-forward mossy fiber pathway in a frequency-dependent manner. A) Simplification of bursting spike trains into two parameter spaces: between burst interval (BI) describing interval between bursts in a spike train, and within BI describing interval between spikes within a burst. B) Example synaptic waveforms of expected mossy fiber EPSCs and IPSCs generated by the short-term plasticity models given short-term plasticity dynamics at three pairs of within and between burst intervals (20ms and 2s, 20ms and 50s, 500ms and 50s respectively). C) Expected short-term plasticity of EPSCs across a wide range of within and between BIs. Light blue, dark blue and black crosses shown in C denote within and between BIs used in the examples shown in B. Data in the presence of CCh not shown since CCh does not change the facilitation of EPSCs. D) Expected short-term plasticity of IPSCs across a wide range of within and between BIs and in the presence and absence of CCh. Pulse ratios for 2^nd^, 4^th^ and 6^th^ pulses compared to the 1^st^ are shown to illustrate change in facilitation across a 6 pulse burst. E) Progression of Excitatory-Inhibitory ratio across the range of within and between BIs and in the presence and absence of CCh. Pulse ratios for 2^nd^, 4^th^ and 6^th^ pulses compared to the 1^st^ are shown to illustrate change in E-I ratio across a 6 pulse burst.

Experimental data shows that mossy fiber EPSCs are exquisitely sensitive to between burst interval with facilitation revealed as between burst interval is decreased. Furthermore, it has been shown that shortening the between burst interval decreases within burst facilitation (Salin et al., 1996). Our simulations replicated this interdependence of between and within burst interval with respect to EPSC facilitation with values closely associated with the experimental data in the literature (Salin et al., 1996; Toth et al., 2000) (Fig. 3B). Since the CCh induced depression of EPSCs was mediated by a reduction in synaptic conductance it did not alter either between or within burst synaptic facilitation.

The situation for inhibitory synaptic transmission was more complex. Over the course of a burst synaptic amplitude facilitation was greatest when between and within burst intervals were largest. As between and within burst intervals reduced the facilitation morphed into a depression towards the end of the burst resulting in limited inhibition at the end of high frequency bursts (Fig. 3C). CCh dramatically reduced both the initial IPSC amplitude and subsequent facilitation within bursts at all between and within burst intervals (Fig. 3C).

We then combined the results from excitatory and inhibitory facilitation to predict EPSC-IPSC amplitude ratios over the course of a burst. In control conditions, excitation dominates over inhibition only after multiple spikes in a burst and when bursts occur at shorter between and within burst intervals (Fig. 3E) (Mori et al., 2004; Torborg et al., 2010; Zucca et al., 2017). However, in the presence of CCh, excitation dominates over inhibition at earlier stimuli within the burst, and over longer between and within burst intervals meaning cholinergic receptor activation allows excitation to dominate over inhibition for a broader range of stimulus patterns.

We next investigated the biophysical effects of the acetylcholine-induced reduction in feed forward inhibitory synaptic transmission at mossy fiber synapses. In particular, the modulation of back-propagating action potentials and EPSPs in CA3 pyramidal cells that are critical for the induction of long-term potentiation (LTP) at recurrent CA3-CA3 synapses and therefore the formation of CA3 ensembles (Brandalise and Gerber, 2014; Guzman et al., 2016; Mishra et al., 2016). Mossy fibers provide powerful excitatory drive to the soma of CA3 pyramidal cells and have been referred to as ‘conditional detonator’ synapses because a single synapse can trigger postsynaptic action potentials in response to high frequency bursts of presynaptic action potentials but not single action potentials (Henze et al., 2002; Sachidhanandam et al., 2009; Vyleta et al., 2016). We hypothesized that feed-forward inhibitory synaptic transmission reduces back-propagating action potentials and mossy fiber evoked EPSPs which will inhibit or prevent the induction of LTP (Tsubokawa and Ross, 1996; Mullner et al., 2015; Wilmes et al., 2016), and that acetylcholine will relieve this inhibition by reducing feed-forward inhibition. To test this, we used a well characterized multi-compartment biophysical model of a CA3 pyramidal cell (Henze et al., 1996; Hemond et al., 2008) with 15 different reconstructed morphologies selected from Neuromorpho.org (Ishizuka et al., 1995; Ascoli et al., 2007). Our model incorporated mossy fiber excitatory synaptic input on the very proximal portion of the apical dendrite and feed-forward inhibitory synapses distributed across both somatic and dendritic compartments (Fig. 4A) (Szabadics and Soltesz, 2009). Membrane potential and the resultant intracellular calcium concentration were simulated across multiple somatic and dendritic compartments of a reconstructed CA3 pyramidal cell incorporating the thin oblique dendrites in stratum radiatum where the majority of CA3-CA3 recurrent synapses are located (Major et al., 1994). With feed-forward inhibition intact, action potentials and EPSPs back-propagate into the principal dendritic shafts without much change in amplitude but are rapidly attenuated on entering the thin oblique dendrites (Fig. 4B and C). This leads to minimal calcium influx through voltage-gated calcium channels at these dendritic sites (Fig. 4B). However, when feed-forward inhibition is reduced by acetylcholine, attenuation of action potentials and EPSPs is greatly reduced allowing substantial calcium influx (Fig. 4B-F). The relief of action potential and EPSP attenuation by acetylcholine was selective for the oblique dendrites in stratum radiatum, was consistent for multiple different CA3 pyramidal cell morphologies (Fig. 4C and Supplementary Fig. 3) and resulted in increases in both the amplitude and probability of action potential backpropagation (Fig. 4D-F). The disinhibition of mossy fiber feed-forward inhibition by acetylcholine has major implications for synaptic plasticity at CA3-CA3 recurrent synapses, since spike timing-dependent plasticity is dependent on the back propagation of action potentials and EPSPs, postsynaptic calcium accumulation and activation of calcium-dependent signalling pathways (Brandalise and Gerber, 2014; Mishra et al., 2016). Our simulations indicate that disinhibition by acetylcholine is a requirement for reliable induction of synaptic plasticity between CA3 pyramidal cells when CA3 ensembles are activated by mossy fiber inputs.

**Figure 4:**
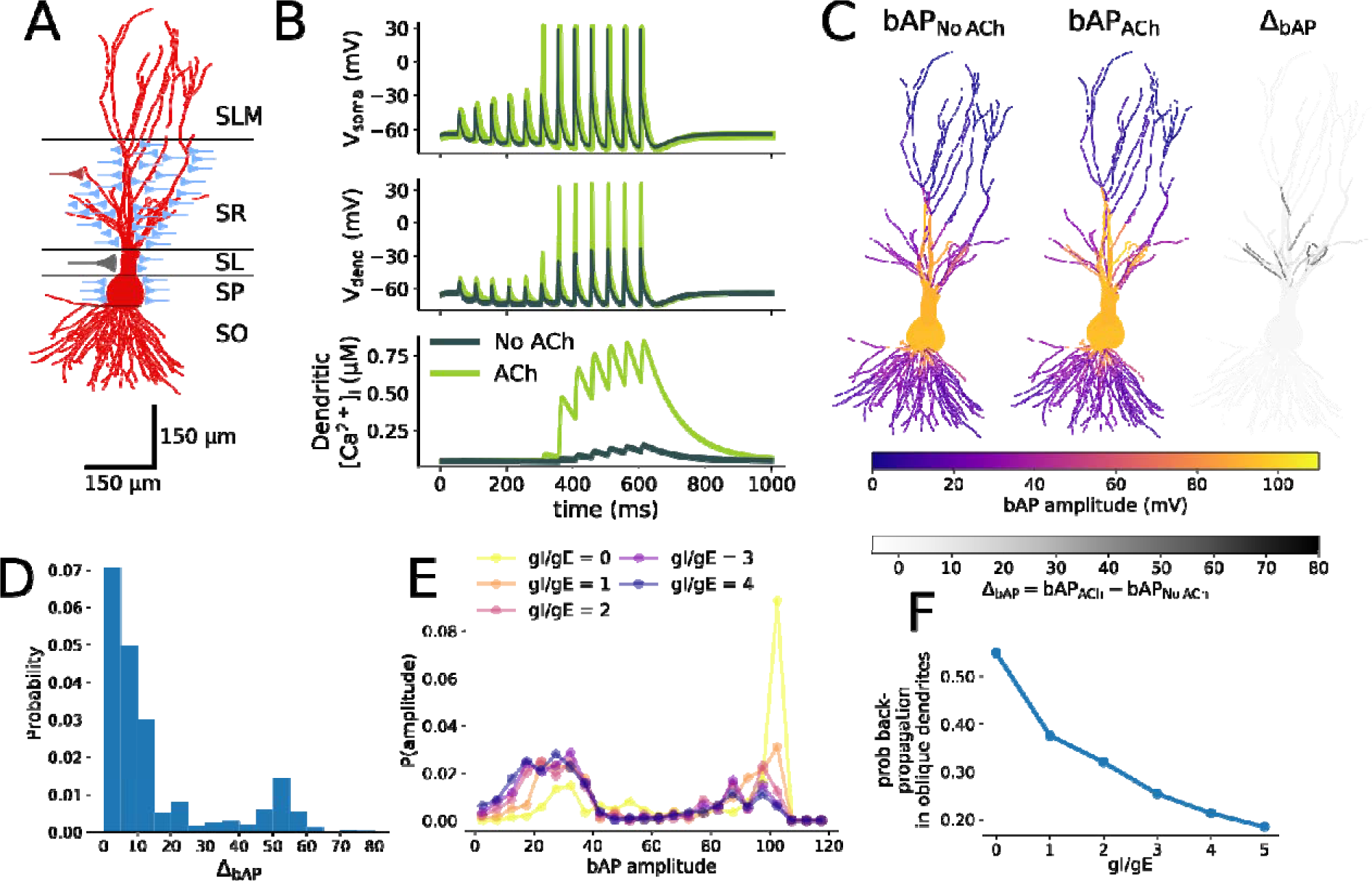
Acetylcholine-mediated disinhibition facilitates back-propagation of EPSPs and action potentials into the dendrites of CA3 pyramidal cells. A) Sketch of CA3 pyramidal cell and positioning within the layers of the hippocampus. Synaptic inputs are shown with location of contact (red - recurrent CA3-CA3 synapse, blue - feed-forward inhibitory synapse, gray - mossy fiber synapse). SLM - stratum lacunosum moleculare, SR - stratum radiatum, SL - stratum lucidum, SP - stratum pyramidale, SO - stratum oriens.B)Example traces produced by the biophysical CA3 neuron model of action potentials generated at the soma from summated mossy fiber EPSPs presented at 20 Hz (top), back-propagation into the radial oblique dendrites (middle), and dendritic calcium influx (bottom), with and without acetylcholine-mediated disinhibition of feed-forward inhibition.C)Back-propagating action potential amplitude before (left) and after (middle) cholinergic modulation, and the difference in amplitude (right) distributed across an example CA3 pyramidal cell. D) Histogram of differences in back-propagating action potential amplitudes with and without acetylcholine disinhibition in stratum radiatum oblique dendritic compartments (< 1 μm diameter) from 15 cells. E) Distribution of back-propagating action potential amplitudes in stratum radiatum oblique dendrites for a range of excitation-inhibition ratios in 15 cells. In our simulations the effect of acetylcholine was modelled as a change in this ratio (the absence of acetylcholine, gI/gE = 3; in the presence of acetylcholine, gI/gE = 1). F) The probability of successful action potential back-propagation (bAP amplitude > 40 mV) in oblique dendrites in 15 cells as a function of excitation-inhibition ratio.

### Ensemble formation in CA3 driven by mossy fiber input

To investigate the effects of acetylcholine on the creation of CA3 ensembles by mossy fiber input we next turned to a spiking network model of CA3. This network was comprised of point neurons with Izhikevich-type dynamics (Izhikevich, 2003) parameterized to reproduce spiking patterns for excitatory CA3 pyramidal cells and inhibitory fast spiking interneurons (Hummos et al., 2014) connected in an all-to-all fashion. Subsets of pyramidal cells were driven by excitatory mossy fiber input with short-term facilitation dictated by the model determined in Fig. 2. CA3-CA3 recurrent synaptic connections were subject to an experimentally determined symmetric spike timing-dependent plasticity (STDP) rule (Fig. 5A) with no short-term plasticity (Mishra et al., 2016). To maintain constant overall network spiking dynamics, the symmetric STDP rule was modified to allow for depression dependent on the postsynaptic firing rate through synaptic scaling. At low firing rates, potentiation is induced with small differences in pre-and post-synaptic spike times. As the postsynaptic firing rates increase, large differences in pre-and post-synaptic spike times cause depression. At a maximum firing rate, no potentiation is possible.

**Figure 5:**
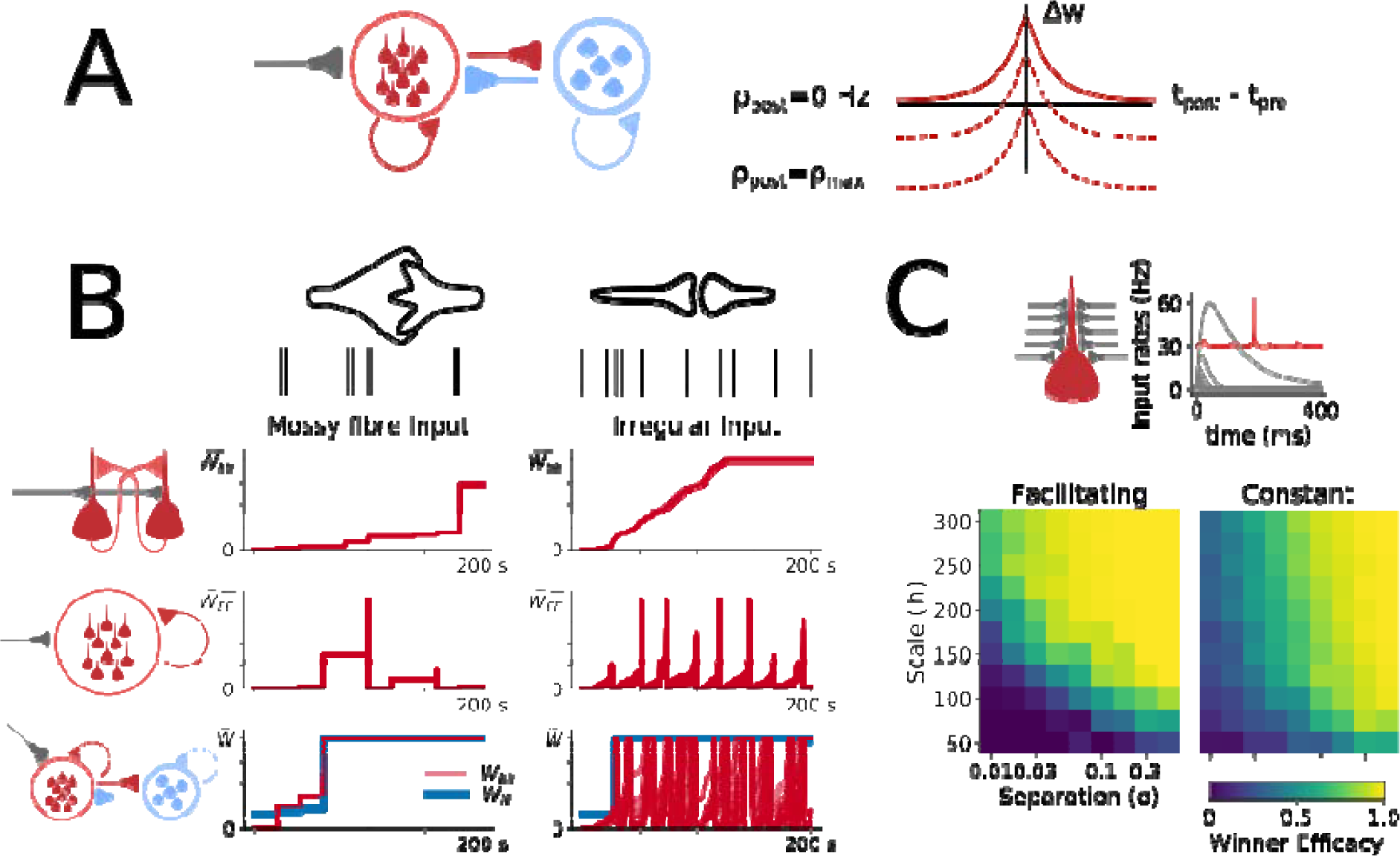
Facilitating mossy fiber inputs generate rapid and stable ensemble formation. A) Schematic showing the network properties of the spiking network model and long-term plasticity rules. Left: A population of excitatory (red) and inhibitory (blue) cells with all-to-all connectivity and mossy fiber input (gray). Right: Recurrent excitatory CA3-CA3 spike timing-dependent plasticity rule with a symmetric window that shifted from potentiation for correlated spiking at low rates (Ppost = 0), to depression for uncorrelated spiking near the maximum postsynaptic firing rate (Ppost = Pmax). B) Comparison of mossy fiber to irregular ‘perforant path’ input. Synaptic weight evolution with time for CA3-CA3 recurrent connections (red) and inhibitory to excitatory connections (blue) for a two cell excitatory population (top), a 10 cell excitatory population (middle) and a population including 10 excitatory and 5 inhibitory cells (bottom). C) Top Left: Network setup - 50 feed-forward inputs to a single granule cell. Top Right: Example of winner-take-all rates and postsynaptic spiking (red) for the population of inputs (gray). This instance shows strong efficacy since postsynaptic spiking occurs once all but one input rates have decreased to near zero. Bottom: Efficacy of winner granule cell to CA3 pyramidal cell coupling with facilitating and constant synaptic input

Within this network we first characterized the speed and stability of ensemble creation where an ensemble was defined as being formed when all synapses between cells within the same ensemble had reached their maximum weight, and all synapses between cells not within the same ensemble had decreased to zero. In addition, the properties of mossy fiber input were studied in comparison with a more generic input reminiscent of perforant path activity during direct information transfer between entorhinal cortex and CA3 to see how they compared in their ability to drive ensemble formation via synchronous spiking in a small population of cells (Fig. 5B). Mossy fiber spike patterns were modelled as a Poisson process with brief (200 ms every 20 seconds) high intensity (50 Hz) firing rates on a very low basal firing rate (0.2 Hz), and were connected to pyramidal cells by a strong facilitating synapse (3.0 nS). This firing pattern represents a strongly separated, sparse firing pattern in a single presynaptic cell, which is expected in dentate gyrus granule cells. Perforant path spike patterns were modelled as a population of presynaptic entorhinal cells in the synchronous irregular state modelled as 120 homogeneous Poisson processes firing at 10 Hz with a correlation coefficient of 0.9 and static, weak synapses fixed at 0.1 nS, which broadly reflects entorhinal activity in a freely behaving rat (Chrobak and Buzsaki, 1998).

We initially built up the network model sequentially to investigate which components were necessary for the speed and stability of ensemble creation. At first, two CA3 pyramidal cells were connected and driven with only excitatory input. For both mossy fiber and perforant path inputs the cells quickly became connected (Fig. 5B, top row). Increasing ensemble size to 10 excitatory cells destabilized the ensemble formation process (Fig. 5B, middle row). The destabilization resulted from unbalanced potentiation of recurrent excitation that caused a large increase in the firing rate leading to strong depression or ‘resetting’ of the synaptic weight with further spiking as a result of synaptic scaling. The addition of 5 feedback inhibitory cells stabilized ensemble formation in the case of mossy fiber input, but not for perforant path input (Fig. 5B, bottom row). This was because the perforant path input provided a colored noise signal amplified by recurrent excitation that caused excitatory cells to fire at high rates too often and feedback inhibition was insufficient to counter this amplification. Mossy fiber input is driven only briefly at sparse intervals, meaning there was little opportunity to exceed target firing rate, and when there was, feedback inhibition was sufficient to contain it. These results show that mossy fiber like sparse inputs and feedback inhibition within the CA3 recurrent network are important for rapid and stable formation of CA3 ensembles.

The advantages of short-term facilitation were also explored. To model how input from multiple granule cells is transmitted to a single pyramidal cell, a single excitatory input layer network to one excitatory cell was constructed (Fig. 5C). Inputs followed an inhomogeneous Poisson process, each of which was connected to the cell with a facilitating mossy fiber synapse, or with a synapse with a fixed conductance at half the maximum weight of the facilitating synapse. Inhomogeneous rates were defined by a winner take all process (Fukai and Tanaka, 1997) with adaptation (see methods). For good transmission of a separated (winner) pattern, the postsynaptic cell should only spike (Fig. 5C; red trace) when the winning input is the only input with a non-zero weight i.e., once the winner has been selected (Fig. 5C; grey traces). The efficacy of transmission is defined as the rate of the winning input *r_N_* at the time of the postsynaptic spike *t_spike_* divided by the sum of rates *r* of all inputs at the time of the postsynaptic spike, i.e.,

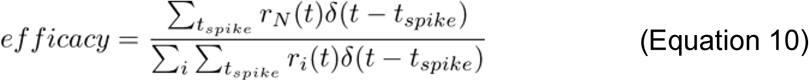

The population of input rates were defined by two parameters: scale and separation, where the scale parameter is a scaling factor that multiplies each rate in the population by the same amount, whereas the separation parameter controls the difference in input to the population of rates, determining how much the winner wins by. Increasing both the scales and separation of input rates produced a sharper transition between no transmission (efficacy = 0) and perfect transmission (efficacy = 1) when synapses were facilitating (Fig. 5C). However, when synaptic conductance was fixed, there was a smoother transition from no transmission to perfect transmission. This indicates that short-term facilitation supports robust transmission of separated patterns, as weakly separated patterns can still be transmitted perfectly. In contrast, weakly separated patterns transmit more ambiguously when input is of fixed conductance. This is because facilitating synapses place very low weight on signals at the start of a winner-take-all process, but much greater weight at the end making spiking more likely once the winner has been chosen, whereas constant synapses place too much weight before a winner has been selected and can cause postsynaptic cells to fire too early.

### Acetylcholine facilitates mossy-fiber driven ensemble formation in CA3

We next examined how acetylcholine affects the CA3 network’s ability to form ensembles. In addition to facilitating plasticity at CA3-CA3 recurrent synapses by control of feed forward inhibition, two well established effects of acetycholine in CA3 are to increase cellular excitability and reduce the overall conductance of CA3-CA3 recurrent synapses by reducing the probability of glutamate release (Hasselmo et al., 1995; Vogt and Regehr, 2001; Dasari and Gulledge, 2011). These cholinergic actions are predicted to increase the number of stored associations within an autoassociative network (Hasselmo et al., 1992; Hasselmo et al., 1995). We implemented these effects of acetylcholine within the spiking network model by depolarizing the resting membrane potential for excitatory and inhibitory cells to -70 mV and -63 mV respectively and halved the CA3-CA3 excitatory synaptic conductance (Hummos et al., 2014). Networks contained 64 excitatory and 16 inhibitory cells with excitatory cells grouped into 8 ensembles consisting of 8 cells each (Fig. 6A). Each cell within a single ensemble received the same mossy fiber input, which followed Poisson process with a low firing rate of 0.2 Hz punctuated by bursts for 250 ms every 20 s at varying frequency and short-term plasticity dynamics dictated by the model determined in Fig. 2. No two ensembles received bursts at the same time and spike timing-dependent plasticity rules were implemented regardless of the presence of acetylcholine.

**Figure 6:**
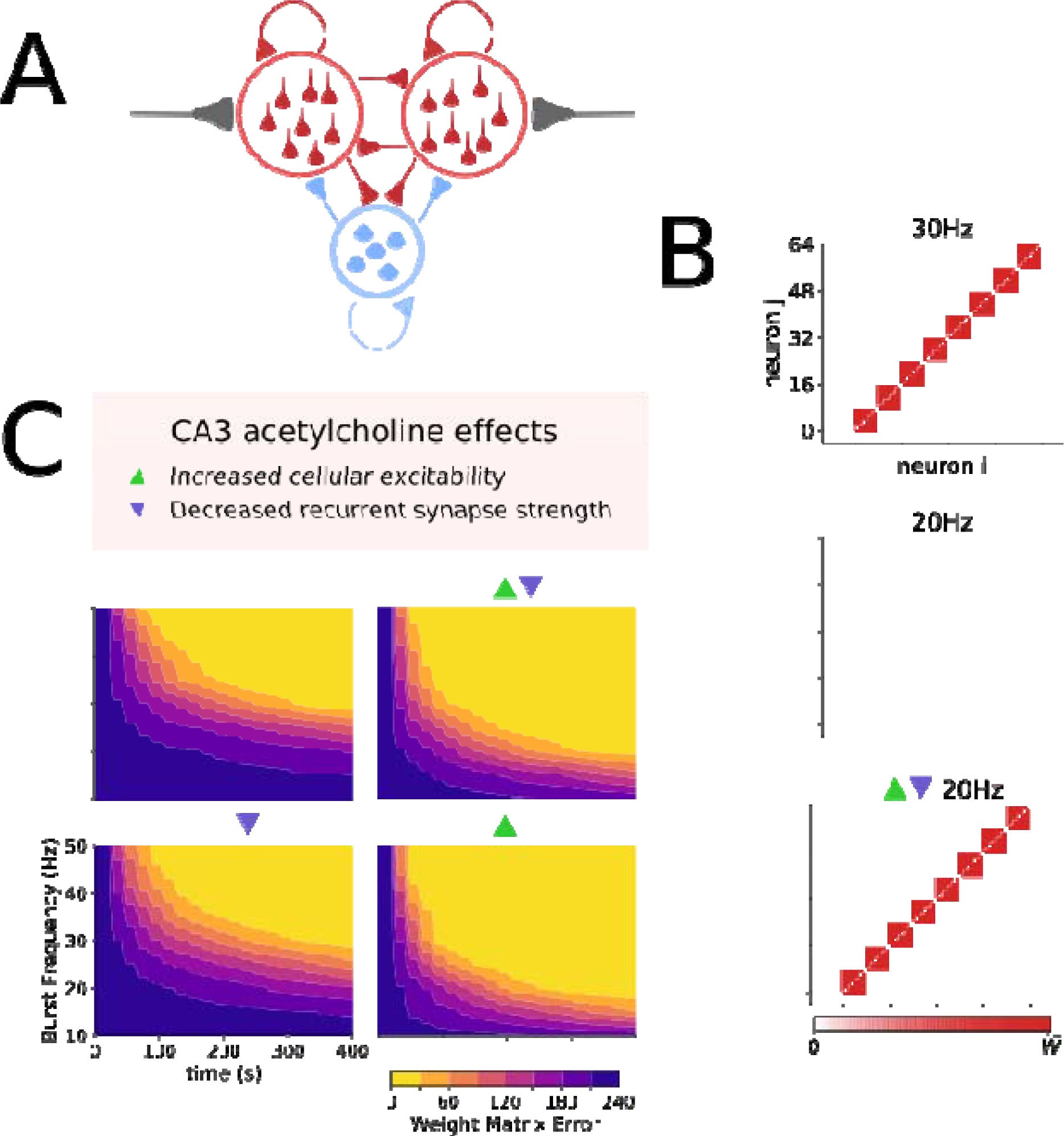
Acetylcholine speeds up ensemble formation and lowers input frequency requirement by increasing cellular excitability. A) Network setup: A population of excitatory and inhibitory cells connected in all-to-all fashion. Subpopulations of excitatory cells receive independent feed-forward input that drives ensemble formation. B) Example weight matrix driven by input with 30 Hz, 20 Hz or 20 Hz bursts with CA3 acetylcholine effects. Stronger weights indicate robust ensemble formation. C) Evolution of ensemble formation illustrated by the weight matrix error reduction over time for different input burst frequencies. Triangles denote which effects of acetylcholine on the CA3 network were included in each set of simulations.

Without acetylcholine, bursts at a frequency of 30 Hz formed discrete ensembles, but these were almost completely abolished when burst frequency was reduced to 20 Hz (Fig. 6B, Supplementary Fig. 5). Remarkably, acetylcholine rescued ensemble formation at the lower burst frequency. To quantify network ensemble formation performance and the impact of acetylcholine in greater detail, we used an error metric (*WME*) defined as the summed absolute difference between target and actual weight matrix 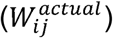, where the target weight matrix 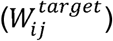 is maximum synaptic weights between cells *i* and *j* in the same ensemble, and zero weight otherwise, i.e.,

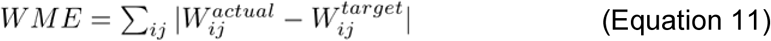

This analysis revealed that acetylcholine lowers the frequency and increases the speed at which ensembles form (Fig. 6C). To test which effects of acetylcholine were critical for these aspects of ensemble formation we removed each parameter change in turn. This revealed that the key factor was the increase in cellular excitability, as removing the parameter changes to cellular excitability abolished the effect of acetylcholine but removing reductions in CA3-CA3 recurrent connections did not (Fig. 6C and Supplementary Fig. 4).

Within the CA3 network multiple often highly overlapping ensembles may be encoded. Theoretically this increases the capacity of information encoding but reduces the fidelity of retrieval with a necessary trade-off between these two parameters. Therefore, we investigated the impact of acetylcholine on the ability of the CA3 network to reliably encode overlapping ensembles. To incorporate overlap between ensembles the total network size was made variable whilst still containing 8 ensembles of 8 cells each and overlap was introduced by having a subset of excitatory cells receive input from two sources rather than one. Overlap was arranged in a ringed fashion such that each ensemble shared a certain number of cells with their adjacent ‘neighbour’ (Fig. 7A). Retrieval was studied by comparing the population rates of each ensemble, with a smaller difference in rates meaning lower discrimination and more difficult retrieval.

**Figure 7:**
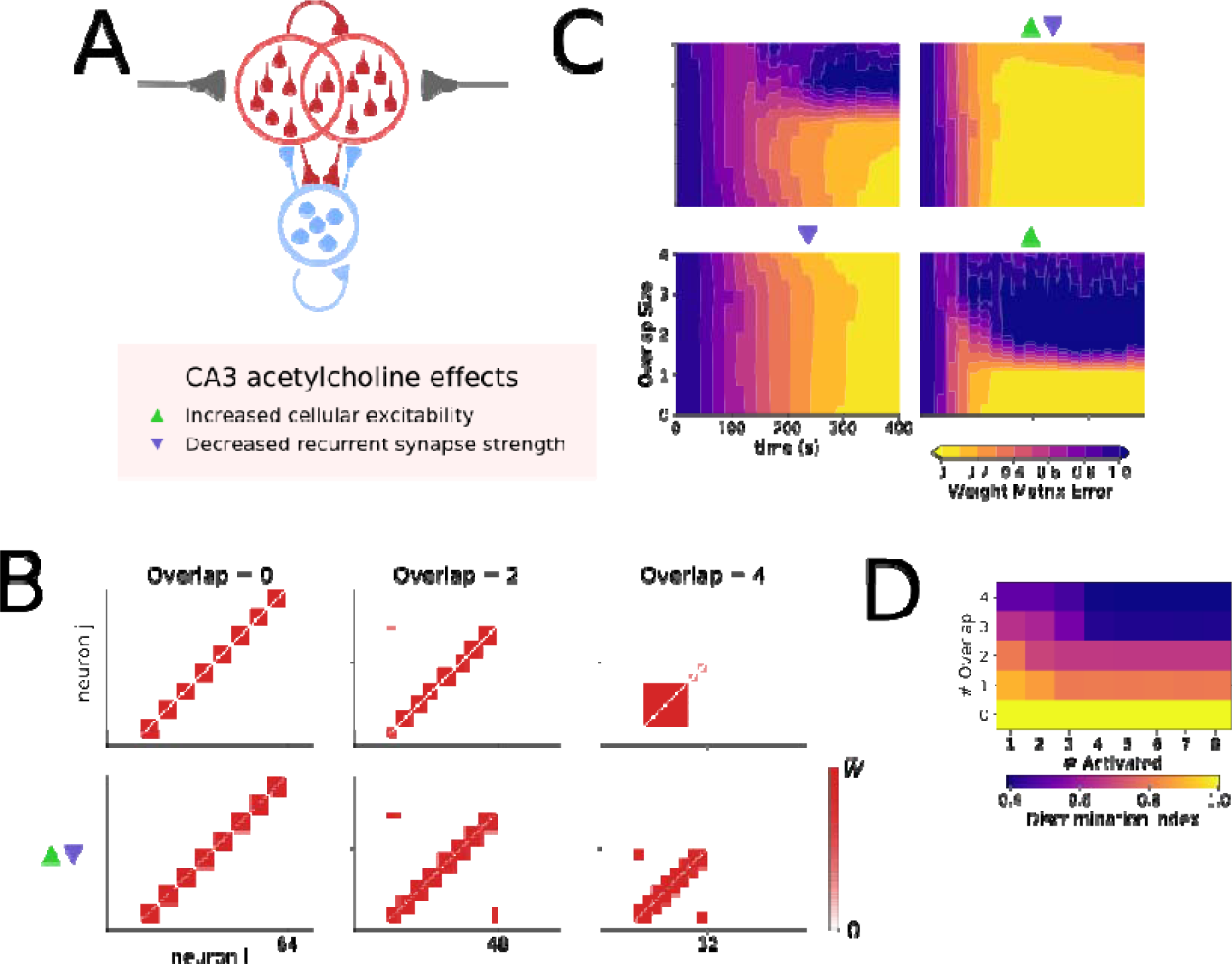
Acetylcholine enables a CA3 network to form stable overlapping ensembles by reducing the strength of recurrent excitatory CA3-CA3 synapses. A)Network setup: A population of excitatory and inhibitory cells connected in all-to-all fashion. Subpopulations of excitatory cells receive overlapping feed-forward input that drives ensemble formation. B) Example weight matrix driven by input with increasing degrees of overlap between ensembles [0, 2, 4 cells]. C) Evolution of ensemble formation illustrated by the weight matrix error over time for different degrees of overlap between ensembles. Triangles denote which effects of acetylcholine were included in the simulation. D) Effect of increasing overlap between ensembles on the ability to discriminate between ensembles defined as the difference in ensemble population spiking rates.

Without acetylcholine present, stable ensembles could be formed when the degree of overlap was low but discrete ensembles could not be formed with levels of overlap >1 (Fig. 7B and C, Supplementary Fig. 6). In contrast, in the presence of acetylcholine the network could safely support an overlap of 3 cells between discrete ensembles (Fig. 7B and C, Supplementary Fig. 6). To test which effects of acetylcholine mediated the enhanced discrimination between overlapping ensembles we removed each parameter change in turn. This revealed that the key factor was the reduction in CA3-CA3 recurrent synapse efficacy since removing this parameter change abolished the ability for acetylcholine to allow greater ensemble overlap. In contrast, removing the increase in cellular excitability only increased the time taken to form stable ensembles without affecting the final degree of overlap supported (Fig. 7C and Supplementary Fig. 4). Interestingly, the increase in stable ensembles with significant overlap was associated with a decrease in the discrimination between ensembles during retrieval of information as more overlap produced less separation of population rates between ensembles (Fig. 7D). Taken together these data indicate that acetylcholine increases the number of discrete ensembles that may be contained within a finite CA3 network but that this comes at a cost of reduced retrieval fidelity.

## Discussion

In this study, we investigated the effects of acetylcholine on the ability of mossy fiber input from dentate granule cells to form ensembles within the CA3 recurrent network. Experimentally, we discovered that acetylcholine dramatically reduces feed-forward inhibition in the mossy fiber pathway whilst having limited effect on mossy fiber excitatory transmission (Figs 1&2). This disinhibition causes a strong positive shift in the Excitatory-Inhibitory balance (Fig. 3) resulting in a greatly enhanced potential for LTP at recurrent CA3-CA3 synapses and therefore ensemble formation (Fig. 4). In addition, in a computational spiking network model, the separate effects of acetylcholine on cellular excitability and basal CA3-CA3 synaptic strength were found respectively to enhance the robustness of ensemble formation (Fig. 6) and the amount of allowable overlap between ensembles (Fig. 7). Together, these findings indicate a central role for acetylcholine in gating and facilitating memory formation in the CA3 network and suggest that cholinergic activity may provide an important salience cue to signal when new ensembles may be formed and therefore which information to encode.

The three separate mechanisms we identify as important for ensemble formation are likely supported by different cholinergic receptors at distinct cellular and subcellular locations. In contrast to the presynaptic actions of cholinergic receptors at other hippocampal synapses (Dasari and Gulledge, 2011), our experimental data show only a small effect of cholinergic agonists at excitatory mossy fiber synapses onto CA3 pyramidal cells indicating a limited direct synaptic modulation by muscarinic or nicotinic receptors mediated by postsynaptic changes (Williams and Johnston, 1990; Vogt and Regehr, 2001; Dickinson et al., 2009). The lack of presynaptic changes indicates no indirect mechanism via enhancement of interneuron spiking and activation of presynaptic GABAB receptors by GABA spillover (Scanziani, 2000; Vogt and Regehr, 2001). For similar reasons, our data do not support a role for presynaptic nicotinic receptors ((Cheng and Yakel, 2014) but see (Vogt and Regehr, 2001)) or an increase in dentate granule cell spike frequency (Vogt and Regehr, 2001) since both would be expected to cause an increase in EPSC amplitude by presynaptic mechanisms. The large depression in feed-forward inhibitory mossy fiber transmission by cholinergic activity most likely results from a combination of an increase in feed-forward interneuron excitability and spike rate, mediated by a combination of muscarinic M1 and M3 receptors and nicotinic receptors (Vogt and Regehr, 2001; Cea-del Rio et al., 2010; Dasari and Gulledge, 2011), coupled with a strong depression of GABA release, mediated by presynaptic M2 receptors present on interneuron terminals (Szabo et al., 2010). This potentially accounts for the observed increase in basal synaptic release to initial stimulation but overall large reduction in synaptic conductance and release over the course of a burst of stimuli seen in our experimental data and short-term plasticity model. Finally, the depression in basal CA3-CA3 recurrent synaptic strength is reported to result from the activation of presynaptic M4 receptors (Dasari and Gulledge, 2011). These mechanistic conclusions predict that M2 muscarinic receptors on interneuron terminals are important for disinhibition of mossy fiber feed-forward inhibition necessary for ensemble formation, M4 muscarinic receptors on CA3-CA3 recurrent axon terminals are important for increasing the amount of permissible overlap between ensembles and M1 muscarinic receptors on CA3 pyramidal cells facilitate the rapid and stable formation of ensembles.

Feed-forward inhibition dominates excitatory transmission in the mossy fiber pathway but unlike other examples of feed-forward inhibition, such as that occurring in the neocortex or CA1 region of the hippocampus, interneurons engaged with mossy fiber feed-forward inhibition target the dendritic compartments of CA3 pyramidal cells as much if not more than the perisomatic areas (Toth et al., 2000; Pelkey et al., 2005; Szabadics and Soltesz, 2009; Torborg et al., 2010). This means that mossy fiber feed-forward inhibition strongly inhibits recurrent CA3-CA3 inputs in stratum radiatum rather than excitatory mossy fiber inputs (Miles et al., 1996; Pouille and Scanziani, 2001). We show that inhibitory input to dendritic domains in stratum radiatum strongly attenuates the back-propagation of action potentials and EPSPs originating from the somatic compartment into the thin radial oblique dendrites where most of the recurrent CA3-CA3 synapses occur (Fig. 4). Since these back-propagating signals are necessary for the induction of LTP at recurrent CA3-CA3 synapses (Brandalise and Gerber, 2014; Mishra et al., 2016), this indicates that mossy fiber feed-forward inhibition is well placed to control the induction of LTP at these synapses. The dominance of mossy fiber feed-forward inhibition may be partially reduced with high frequency burst stimulation where feed-forward inhibition does not facilitate as strongly as excitation enabling excitation to dominate (Mori et al., 2004) (Fig. 3) although this is not the case during development of the mossy fiber pathway (Torborg et al., 2010). Here, the remarkable finding is that cholinergic activation reduces mossy fiber feed forward inhibition by >70% (Fig. 1&2) removing the attenuation of back-propagating action potentials and EPSPs (Fig. 4), which therefore enables LTP at CA3-CA3 recurrent synapses.

Further investigation revealed that not only does acetylcholine enable ensemble formation by reducing mossy fiber feed-forward inhibition, but it also alters the properties of the CA3 network to allow ensemble formation to occur rapidly and robustly with a high degree of overlap between ensembles. This supports findings in similar models of piriform cortex and CA3 attractor networks (Hasselmo et al., 1992; Hasselmo et al., 1995). Enhancing cellular excitability within an attractor network such as the recurrent CA3 network increases the speed and robustness of synaptic plasticity due to increased spiking during ensemble activity (Fig. 6). For stable ensemble formation and network configuration the increased excitability must be regulated by feedback inhibition (Fig. 5). Theoretically, the reduction in CA3-CA3 synaptic efficacy caused by acetylcholine (Hasselmo et al., 1995) might be predicted to reduce the efficiency of ensemble formation but our results show this is not the case (Fig. 6) likely because of reduced interference between ensembles (Hasselmo et al., 1992). Furthermore, we found that this effect of acetylcholine enabled a greater overlap between ensembles whilst still maintaining their discrete identity (Fig. 7). This is important for a couple of reasons: i) it allows an increased density of discrete ensembles to be encoded which in a finite network will increase its capacity to store information, and ii) an increase in overlap between ensembles has been suggested to facilitate memory consolidation and generalization during reactivation of ensembles that occurs during sleep (O’Donnell and Sejnowski, 2014). Interestingly, our model used a method to limit synaptic strengthening based on recently described STDP rules (Mishra et al., 2016) coupled with a rate-based scaling factor, whereas previous models of similar autoassociative networks have used an LTP only rule with a saturation function (Hasselmo et al., 1992; Hasselmo et al., 1995). Remarkably, both these methods produced very similar outcomes indicating that the effects of acetylcholine on the rate and degree of overlap for ensemble creation are independent of different plasticity rules. A more important factor may be non-linear dendritic conductances which increase the storage capacity for similar or overlapping memories within the CA3 network (Kaifosh and Losonczy, 2016). Future studies may determine how the mechanisms engaged by acetylcholine and non-linear conductances interact and combine within the hippocampal CA3 network.

A core symptom of Alzheimer’s disease is deteriorating episodic memory, which may be ameliorated by treatment with cholinesterase inhibitors. However, the mechanisms by which increasing the availability of acetylcholine in the brain provides this cognitive benefit remain obscure. At the behavioral level, our findings predict that cholinesterase treatment facilitates the formation of memory ensembles within the hippocampus and increases the storage capacity for separate memory representations. It is widely reported that cholinesterase inhibitors provide cognitive enhancement (McGleenon et al., 1999) but the specific cognitive domains affected are less well characterized. At a network level, our findings predict that interventions to deprive the hippocampus of cholinergic innervation will prevent the update of ensemble configurations in CA3 (Atri et al., 2004) and, furthermore, that stimulation of acetylcholine release at specific locations will bias ensemble formation towards the incorporation of place cells representing those locations. Manipulations of acetylcholine release in the hippocampus have largely focussed on the effects on oscillatory activity where acetylcholine has been found to promote theta activity and suppress sharp wave ripples (Vandecasteele et al., 2014). However, in support of our predictions, cholinergic activation has also been found to increase the number of neurons incorporated into ensembles measured by their activity during sharp wave ripples (Zylla et al., 2013).

Acetylcholine release within the central nervous system has classically been portrayed as a signal for arousal and attention and is strongly associated with learning (McGaughy et al., 2000; Parikh et al., 2007; Hasselmo and Sarter, 2011). This model has been adapted to propose that acetylcholine is released in response to environments where the outcome is uncertain or not as predicted by internal representations (Yu and Dayan, 2005). In such a scenario new information needs to be incorporated into the internal representation to make the environment more familiar and the outcomes more predictable (Hasselmo and Sarter, 2011). Acetylcholine supports this process by enabling the updating of internal representations of the environment (episodic memories). Our data support such a model where acetylcholine reconfigures the dentate gyrus and CA3 microcircuit to enable the formation of memory ensembles within the recurrent CA3 network.

## Materials and Methods

### Ethical approval

All experiments were performed in accordance with the UK Animal Scientific Procedures Act (1986) and local guidance from the Home Office Licensing Team at the University of Bristol. The protocol was approved by the Animal Welfare and Ethics Review Board at the University of Bristol.

### Slice Preparation

500 μm thick transverse slices of the hippocampus were prepared from 4-6 week old C57/BL6 mice. After cervical dislocation, brains were removed and submerged in ice-cold cutting solution saturated with oxygen (in mM: 85 NaCl, 75 Sucrose, 2.5 KCl, 25 Glucose, 1.25 NaH2PO4, 4 MgCl2, 0.5 CaCl2, 24 NaHCO3). Each hippocampus was dissected out and mounted onto a cube of agar then glued to the slicing plate such that hippocampi were positioned vertically and cut using a Leica VT1200 vibratome. Slices were then transferred to a holding chamber with oxygenated aCSF (in mM: 119 NaCl, 2.5 KCl, 11 Glucose, 1 NaH2PO4, 26.5 NaHCO3, 1.3 MgSO4, 2.5 CaCl2), incubated for 30 mins at 35°C and left to rest for a further 30 minutes - 5 hours at room temperature.

### Electrophysiology

Slices recordings were made in a submerged recording chamber at 33-35°C. CA3 pyramidal cells were visually identified using infra-red differential interference contrast on an Olympus BX-51W1 microscope. Recording pipettes with resistance 2-4 MΩ were pulled from borosilicate filamented glass capillaries and filled with a caesium-based intracellular solution (in mM: 130 CsMeSO3, 4 NaCl, 10 HEPES, 0.5 EGTA, 10 TEA, 1 QX-314 chloride, 2 MgATP, 0.5 NaGTP). Series resistances were continuously monitored and recordings discarded if series resistance > 25 MΩ or changed by >50%. Recordings were collected using a Multiclamp 700A amplifier (Molecular Devices) filtered at 4 kHz and sampled at 10 or 25 kHz using Signal or Spike2 acquisition software, and a CED Power 1401 data acquisition board.

Postsynaptic currents were evoked by placing monopolar stimulation electrodes in the granular layer of the dentate gyrus and applying 200μs pulses driven by a Digitimer DS2A Isolated stimulator. Stimulating the granular layer avoids the risk of contamination from perforant path inputs, associational/commissurals, or monosynaptic inhibitory synapses. Excitatory currents were obtained by holding the cell in voltage-clamp mode at -70 mV, while inhibitory currents were obtained by holding at +10 mV. Candidate mossy fiber driven responses were chosen based on their latency (2-3 ms for excitatory, 5-10 ms for inhibitory). Excitatory currents were also selected based on kinetics (< 1 ms 20-80% rise time) and short-term facilitation in response to 4 pulses at 20 Hz. Stimulation strength was calibrated to the minimum strength that evoked a response (minimal stimulation). Excitatory and inhibitory responses were blocked > 80% by 1 μM DCG-IV in 100% cases (Fig. 1; n=9 MF-EPSCs, n=6 MF-IPSCs), therefore it can be inferred that this approach reliably stimulated mossy fibers (Kamiya et al., 1996). Carbachol (CCh) was bath applied for 5-10 minutes before measuring effect on synaptic transmission. Effects of carbachol were normalised to control values taken in first 3 minutes of the experiment.

### Short-Term Plasticity Model

The Tsodyks-Markram model (Tsodyks and Markram, 1997; Hennig, 2013) was adopted due to its widespread use, simplicity, and relation to biophysics. This model captures pre synaptic release dynamics with two variables, a facilitating process *f* and a depressing process d, that represent ‘resources’ available to drive synaptic transmission. These evolve as follows:

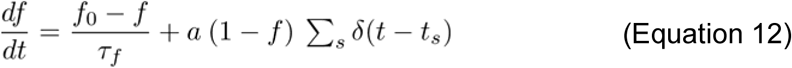

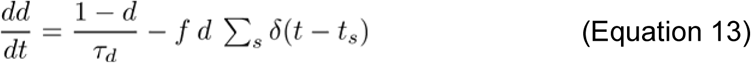

The dynamics of *f* are governed by three parameters: *f_0_* (baseline value of *f*), *τ_f_* (decay time constant of *f*), and *a* (increment scaling factor with incoming spike at time *t_s_*). This variable loosely represents the build-up of free calcium ions in the presynaptic terminal that triggers exocytosis of neurotransmitter-containing vesicles into the synaptic cleft. Dynamics of *d* is governed by a single parameter *τ_d_*, and loosely represents the availability of docked vesicles at release sites. Since these variables are bounded between 0 and 1, the convert into a conductance amplitude these are multiplied by a conductance scaling factor *g,* i.e.,

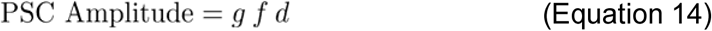

This basic model can be extended to more complex facilitation models for example by making parameters *f_0_* and *a* time dependent (Hennig, 2013), i.e.,

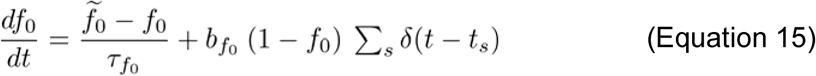

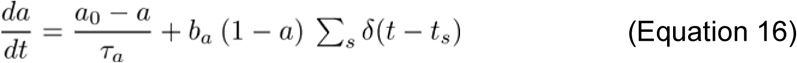

where 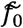 a baseline for *f_0_*, *a_0_* is the baseline for a, *τ_f_0__* is the time constant for *f_0_, T_a_* is the time constant for a, *b_fo_* is the increment scaling factor for *f_0_*, and *b_a_* is the increment scaling factor for a. It can also be reduced by making *f* or *d* constant. Since the mossy fiber synapse is well known for its large pool of readily releasable vesicles that can be quickly replenished, a simple reduction is to keep *d* constant at 1, which is true when *T_d_* << min(/*S*/). Further complexity can be incorporated by allowing multiple independent depressing variables, or by having a time dependent scaling factor

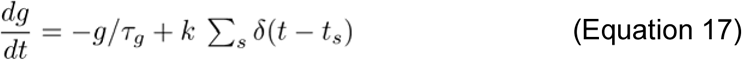

where *τ_g_* is the time constant for short-term changes in conductance, and *k* is an increment scaling factor that can take positive values for facilitation, or negative values for depression.

### Naturalistic stimulation patterns

Previous research has shown that short-term facilitation models are difficult to constrain with responses evoked by regular stimulation protocols (Costa et al., 2013), and that irregular or naturalistic stimulus trains allowed much better fits to data due to sampling across a broader range of inter-stimulus intervals (ISI) (Gundlfinger et al., 2007; Costa et al., 2013). Dentate gyrus granule cells in vivo have been shown to have bimodal ISI distributions, with long periods of quiescence punctuated by short bursts of action potential firing (Jung and McNaughton, 1993; Mistry et al., 2011). This bimodal ISI distribution was modelled as a doubly stochastic Cox process to allow generation of stimuli resembling natural spike patterns (Dayan and Abbott, 2001). Each Cox process *i* is defined by a rate parameter *λ_ii_,* and a refractoriness parameter *σ_i_*. These two processes are then mixed with responsibility *π* i.e.,

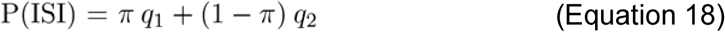

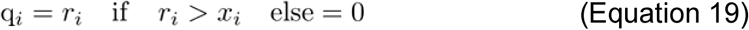

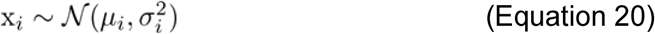

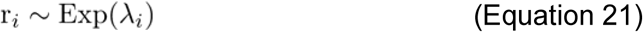

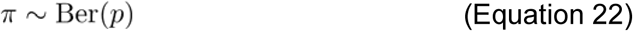

In brief, ISIs were generated by sampling from two exponential distributions with parameters *λ_i_,* which were rejected if they were less than a sample from a normal distribution with standard deviation *σ_i_* These candidate ISIs were then accepted according to a Bernoulli distribution with probability *П* These parameters were set as *λ_1_* = 3.0, *λ_2_* = 0.25, *σ_1_* = 0.12, *σ_2_* = 0.01, *π*= 0.55. Ninety-nine ISIs were sampled to provide 100 spike times for a stimulation protocol lasting 525 seconds.

### Model Fitting

The Bayesian parameter inference procedure used by (Costa et al., 2013) was used to fit parameters to the model. Each model was converted into an iterative form, integrating *f* and *d* over each ISI Δ*t_s_* between the *n^th^* and *n+1^th^* spike to output a normalised post-synaptic conductance amplitude for the *n^th^* spike in the sequence, i.e.,

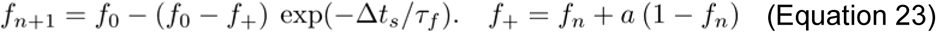

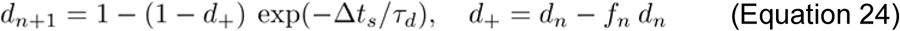

from the original Tsodyks-Markram formalism, however for more complex models in which *a* and *f_0_* are time dependent

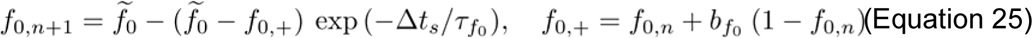

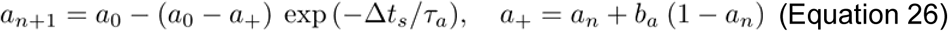

Additionally, when *f_0_* is time-dependent, Equation 22 becomes

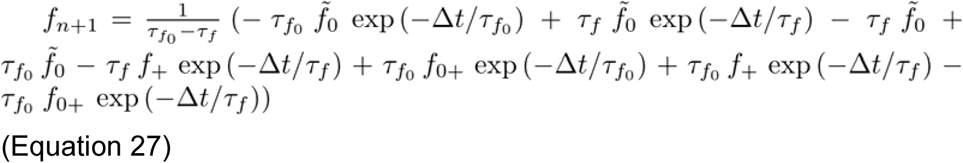

These were then compared to PSC amplitude estimated by the difference between response peak and a baseline taken just before stimulus onset.

Since these equations are deterministic, a likelihood model was constructed where the amplitude was used as parameters for a normal distribution, i.e.,

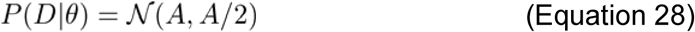

where *A* is PSC amplitude (Equation 14).

Exponential priors were used for conductance scaling parameters, beta priors for baseline and increment parameters, and uniform priors for time constants. This was to bias conductances towards smaller values and keep baselines low as would be expected from facilitating synapses. Posterior distributions were estimated using Markov Chain Monte Carlo sampling via the Metropolis-Hastings algorithm using the pymc python module.

Model selection was conducted using AIC and BIC weights (Wagenmakers and Farrell, 2004), a transformation of AIC and BIC values into a probability space, with best model having the highest weight. This made it possible to compare the best fitting model over the population when fitted for each individual sample. For the information criterion of a model for a given sample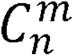 this is defined as

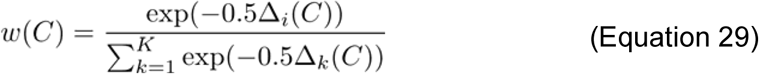

where

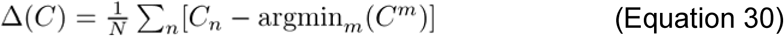

Goodness-of-fit for the best fitting model was assessed by estimating Bayesian poster-predictive *p*-values, where samples are drawn from the posterior-predictive distribution and discrepancies *D*(*x*|*θ*) to expected values *e* from the model *x_sim_* and to the data *x_obs_* are compared (Gelman et al., 1996), i.e.,

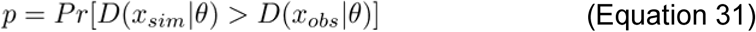

where

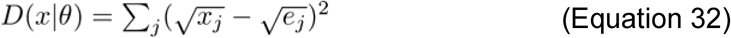

If discrepancies were similar, i.e., *p* >0.025 or < 0.975, then the model is assessed to fit well. This quantifies how easy it is to discriminate between posterior samples and actual data.

### Compartmental Modelling

A 943 compartment reconstruction of a CA3 pyramidal cell with active dendrites (Henze et al., 1996; Hemond et al., 2008) was used to study whether mossy fiber feed-forward inhibition could regulate action potential backpropagation, which would be necessary for feed-forward inhibition to regulate plasticity between recurrent synapses. Simulations were carried out using NEURON.

Active conductances included voltage-gated sodium (Na_V_), voltage-activated potassium conductance including delayed rectifier (K_DR_), M-current (K_M_), fast-inactivating A-type (K_A_), calcium conductances including N-type (Ca_N_), T-type (Ca_T_), and L-type (Ca_L_), calcium-activated potassium conductances (K_C_ and K_AHP_). Calcium extrusion was modelled as a 100ms decay to a resting Ca^2+^ of 50 nM. Channel kinetics were similar to those used in other hippocampal pyramidal neuron models (Migliore et al., 1999).

Somatic compartments contained all conductances, dendritic compartments contained all except K_M_, and the axonal compartment contained only Na_V_, K_DR_, and K_A_. For action potential generation, sodium conductance was five times higher in the axon than the rest of the neuron. Conductances were set as (in μS/cm^2^): gNa_V_ = 0.022, gK_DR_ = 0.005, gK_M_ = 0.017, gK_A_ = 0.02, gCa_N_ = 0.00001, gCa_T_ = 0.00001, gCa_L_ = 0.00001, gK_C_ = 0.00005, gK_AHP_ = 0.0001.

Dendritic compartments along the apical dendrite were subdivided according to distance (in microns from soma) into those within stratum lucidum (≤ 150), stratum radiatum (>150 or ≤400), and stratum lacunosum moleculare (>400). Mossy fiber synapses were targeted towards compartments in stratum lucidum. Feed-forward inhibition was targeted towards somatic compartments (50%) and dendritic compartments in stratum lucidum and stratum radiatum (50%) reflecting the diversity of interneuron subtypes and their targets (Szabadics and Soltesz, 2009).Synaptic input was modelled using bi-exponential kinetics and short-term plasticity dynamics fit to experimental data. The effect of carbachol was modelled as a three-fold reduction in feed-forward inhibitory conductance.

### CA3 Network Modelling

The hippocampal CA3 region was modelled as a small all-to-all recurrent network comprised of excitatory and inhibitory point neurons with adaptive quadratic-integrate-and-fire dynamics with parameters to reflect the firing patterns in response to current injection of CA3 pyramidal cells and fast-spiking basket cells respectively (Izhikevich, 2003; Hummos et al., 2014). Continuous membrane dynamics for neuron *i* is described by two equations:

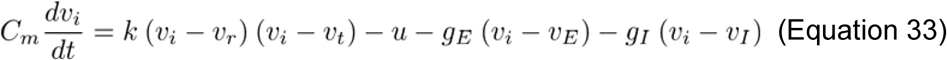

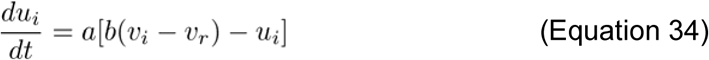

where *v_i_* describes membrane potential, and *u_i_* is a slow adaptation variable. Parameters for excitatory cells were: *C_m_* = 24 pF, *k* = 1.5 pA/mV^2^, *a*= 10 Hz, *b* = 2 nS, *c* = -63 mV, *d* = 60 pA, *v_r_* = -75, *v_t_* = -58 mV, *v_peak_* = 29 mV. Parameters for inhibitory cells were: *C_m_* = 16 pF, *k* = 1.5 nS/mV, *a* = 900 Hz, *b* = 2 nS, *c* = -80 mV, *d* = 400 pA, *v_r_* = -65 mV, *v_t_* = -50 mV, *v_peak_* = ^28 mV^.

Synaptic reversal potentials were set as *v_E_* = 10 mV, and *v_I_* = -80 mV. When *v_i_,* ≥ *v_peak_, v_i_,* and *u_i_* were reset to

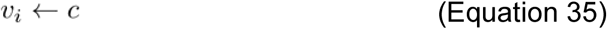

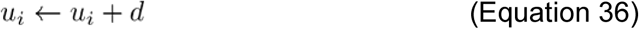

Excitatory and inhibitory cells were connected through four types of synapses: excitatory to excitatory cell (EE) synapses, excitatory to inhibitory cell synapses (EI), inhibitory to excitatory cell synapses (IE), and inhibitory to inhibitory cell (II) synapses. Kinetics were modelled as exponential synapses such that the synaptic conductance *g_syn_* evolved according to:

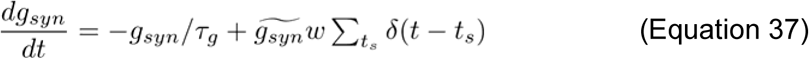

where 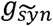 the maximum synaptic conductance, and *τ_g_* = 10 ms for EE and EI synapses and 20 ms for IE and II synapses. Maximum excitatory synapse conductance was 0.5 nS, and 1.0 nS for inhibitory synapses. Cholinergic modulation was implemented by changing a subset of network parameters (See Table 1).

**Table 1.**
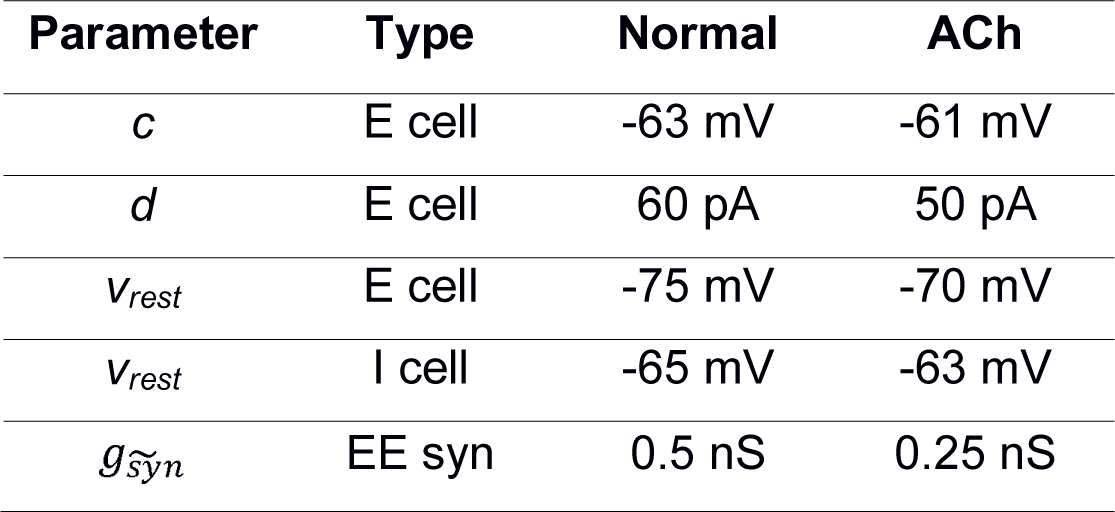
CA3 network parameter changes to model cholinergic modulation of CA3

EE and IE synapses were subject to spike timing-dependent plasticity of the form

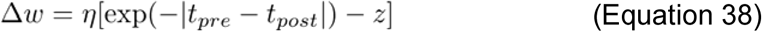

where *η* is a learning rate, and *z* is a scaling factor. This provides a symmetric STDP rule used previously as a homeostatic means to balance excitation and inhibition through inhibitory plasticity, (Vogels et al., 2011) and has also recently been shown to operate at CA3 associated commissural synapses (Mishra et al., 2016). In the case of IE synapses, *η* and *z* are fixed, however in the case of EE synapses these evolve according to

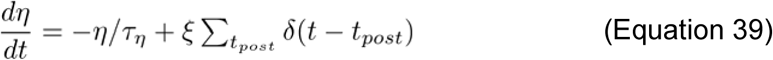

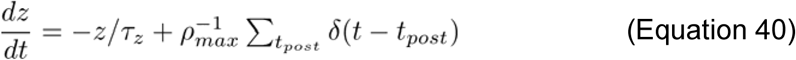

This makes *η* and *z* track the postsynaptic firing rate for two different purposes, and on different time scales since *τ_η_* = 100 ms, and *τ_z_* = 1 second; *η* becomes a burst detector that increases the learning rate by a factor *ξ* meaning STDP requires multiple post-synaptic spikes to be activated, and *z* scales STDP such that the postsynaptic firing rate reaches a maximum *ρ_max_,* more pre-and post-synaptic spike pairs cause depression, preventing STDP from inducing unrealistically high firing rates in an excitatory recurrent network. Additionally, STDP was bounded between 0 and the maximum conductance of the synapse.

Synaptic input to cells was comprised of recurrent and feed-forward inputs, i.e.,

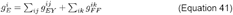

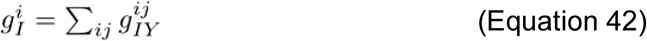

where g_FF_ is the conductance of feed-forward input, g_EY_ is excitatory recurrent input, and g_IY_ is recurrent inhibition. Feed-forward input was given only to excitatory cells. CA3 Network architecture and feed-forward dynamics varied according to each simulation.

Granule cell spiking was modelled as an inhomegenous Poisson process. For most simulations, it was modelled low spike rate of 0.2 Hz, punctuated by jumps to a much higher burst frequency every 20 seconds for 250 ms. In one set of simulations, a small granule cell population (*N* = 50) was modelled as a set of Poisson processes rates defined by winner-take-all dynamics with adaptation (Fukai and Tanaka, 1997)

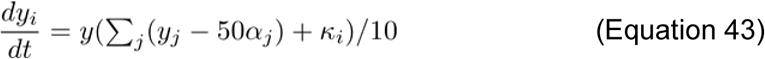

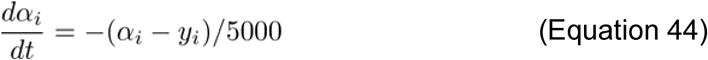

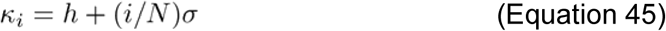

where *y* are granule cell rates following winner-take-all dynamics, *α* is rate-adaptation taking place over a longer time scale, and *κ* is the input to each granule cell determined by *h* and *σ* which define the scale and separation of rates respectively, where the scale parameter is a scaling factor that multiples each rate in the population by the same amount, whereas the separation parameter controls the difference in input to the population of rates, determining how much the winner wins by. The winner granule cell receives input *κ* = *h* + *σ*

Network retrieval performance is measured by a discrimination index that imagines that ensemble population rates are read by a downstream neuron. The greater the difference in ensemble population rates, the easier it is to differentiate between ensembles and retrieval is more precise. The discrimination index *D* is defined as

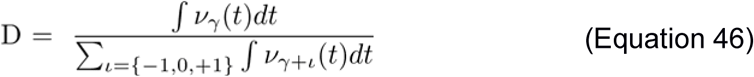

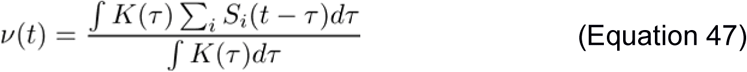

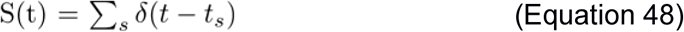

where *v_Y_(t*) is the population rate *v* of ensemble *γ, S_i_(t)* is the spike train *S_i_(t*) of neuron *i, K(t*) is a kernel averaging the spike train over a defined window, and 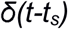 is the delta function modelling a spike at time *t_s_*. This discrimination index essentially calculates the ratio *signal/(signal + noise*) where the signal is the population rate of the ensemble representing the memory being retrieved, and the noise is the population rate of ensembles representing memories that should not be retrieved and are interfering with the retrieval process. As such, when *D* is smaller the interference from neighbouring neurons is higher.

## Acknowledgements

We thank members of the Mellor lab for helpful discussion and C. O’Donnell and M. Ashby for comments on previous versions of the manuscript. LYP, CC and JRM funded by the Wellcome Trust. The project was also supported by an IBRO & Simons fund through Grant ID # isiCNI2017. KTA gratefully acknowledges the financial support of the EPSRC via grant EP/N014391/1. The authors confirm no conflict of interest.

## Author Contributions

Conceptualization, L.Y.P. and J.R.M.; Methodology, L.Y.P., K.T.-A. and C.C.; Investigation and Analysis, L.Y.P.; Writing, L.Y.P., K. T.-A., C.C. and J.R.M.; Supervision, K.T.-A., C.C. and J.R.M.

---

**Supplementary Figure 1:**
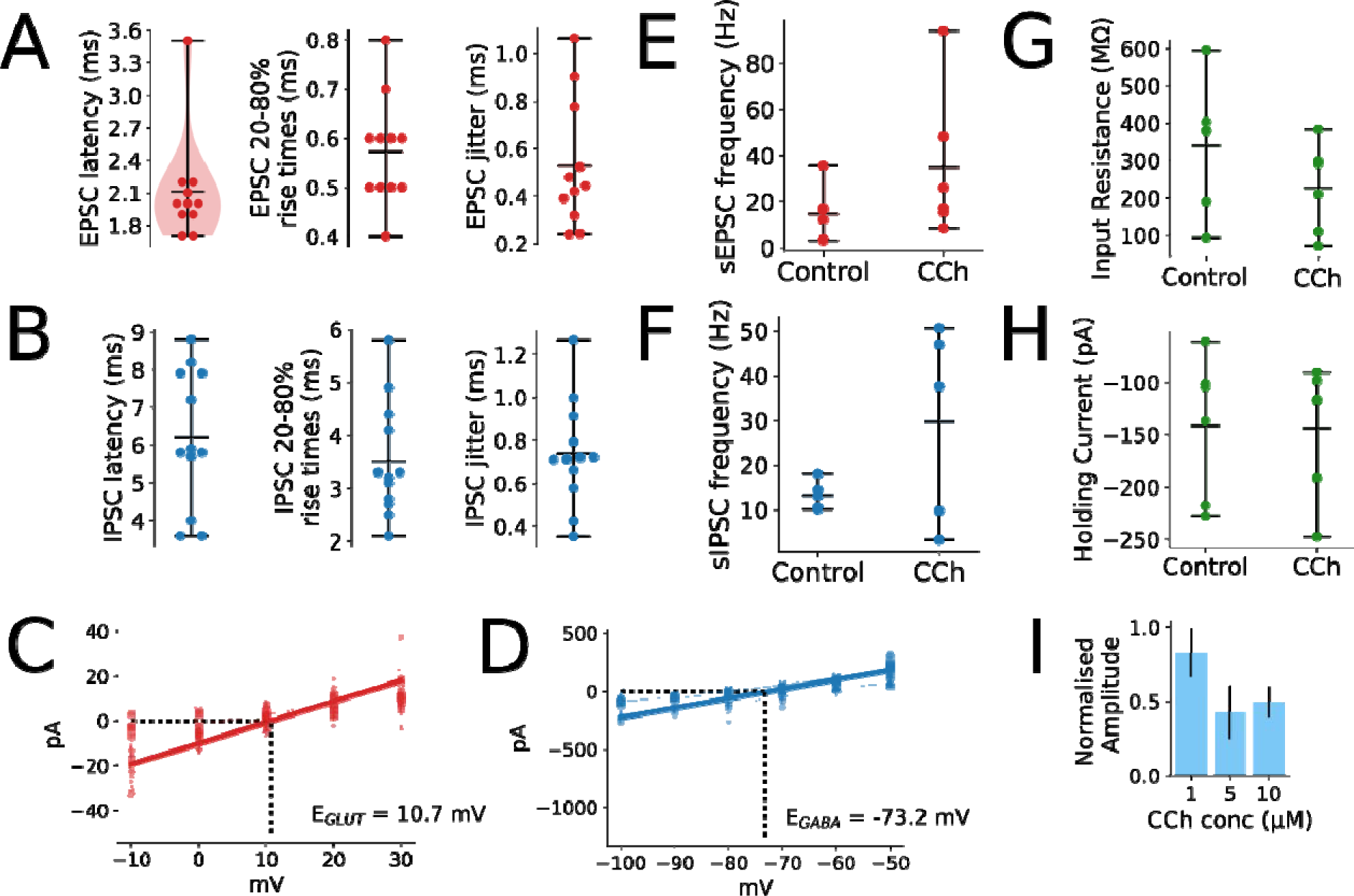
A-B) Latency, rise times, and jitter of mossy fiber driven EPSCs (n = 11) (A) and IPSCs (n = 12) (B). C-D) Reversal potential estimation of glutamatergic (n = 5) (C) and GABAergic (n = 6) (D) transmission at CA3 pyramidal cells. E-F) Spontaneous EPSC (n = 6) (E) and IPSC (n = 5) (F) frequency recorded before and after carbachol application. G-H) CA3 pyramidal cell input resistance (n = 5) (G) and holding current at -70 mV (n = 5) (H) before and after carbachol application. I) Dose-response of carbachol effect on IPSC amplitudes (n = 3).

**Supplementary Figure 2:**
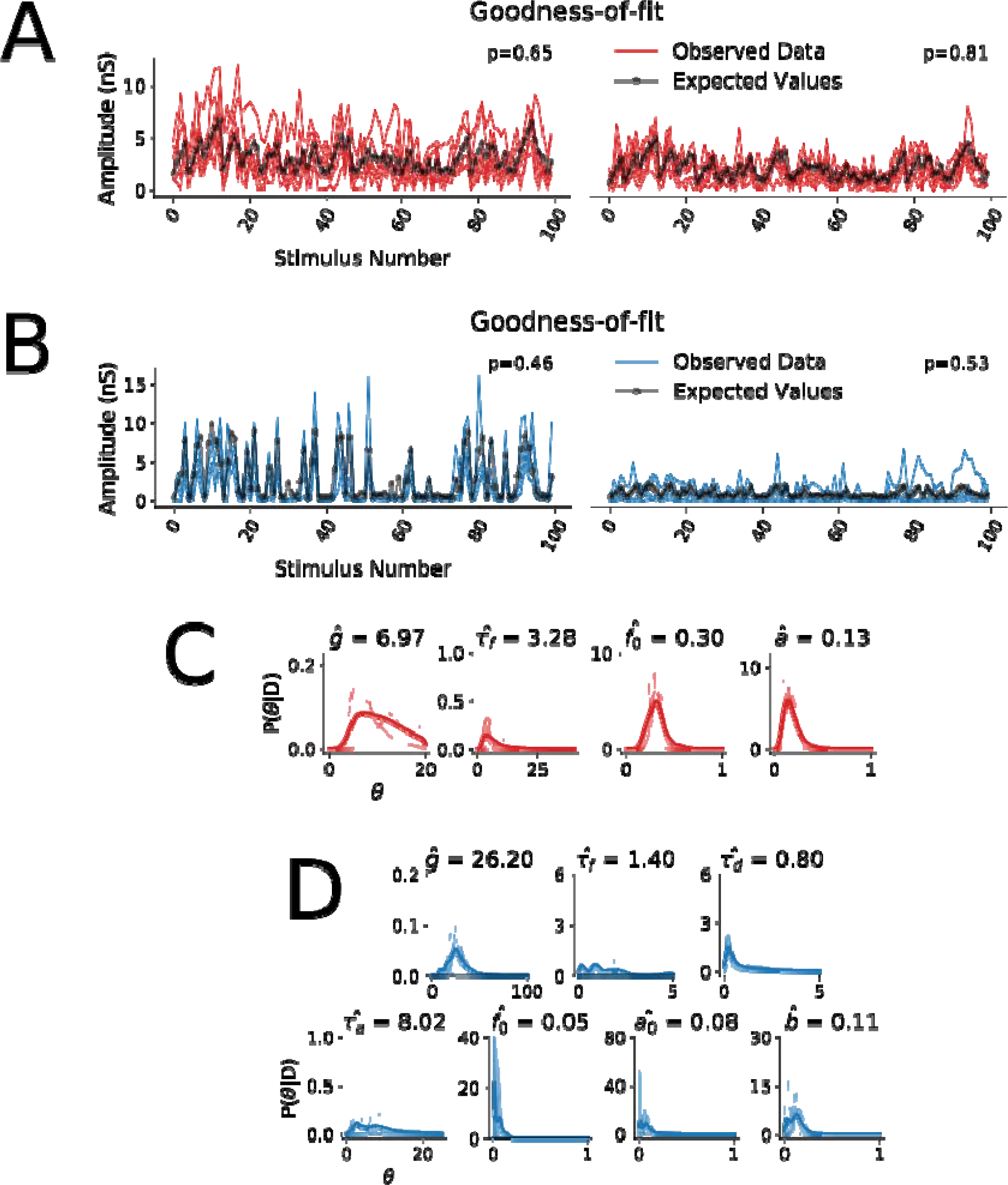
A-B) Goodness-of-fit for EPSC (A) and IPSC (B) short-term plasticity models assessed by Bayesian posterior predictive p-values. Plots show the observed data for all experiments (EPSCs and IPSCs respectively) together with the expected values from the model before and after the application of 5 μM carbachol. p-values close to 0.5 indicate best fit. C-D) Posterior distributions for parameters of best fitting models given data for EPSCs (C) and IPSCs (D).

**Supplementary Figure 3:**
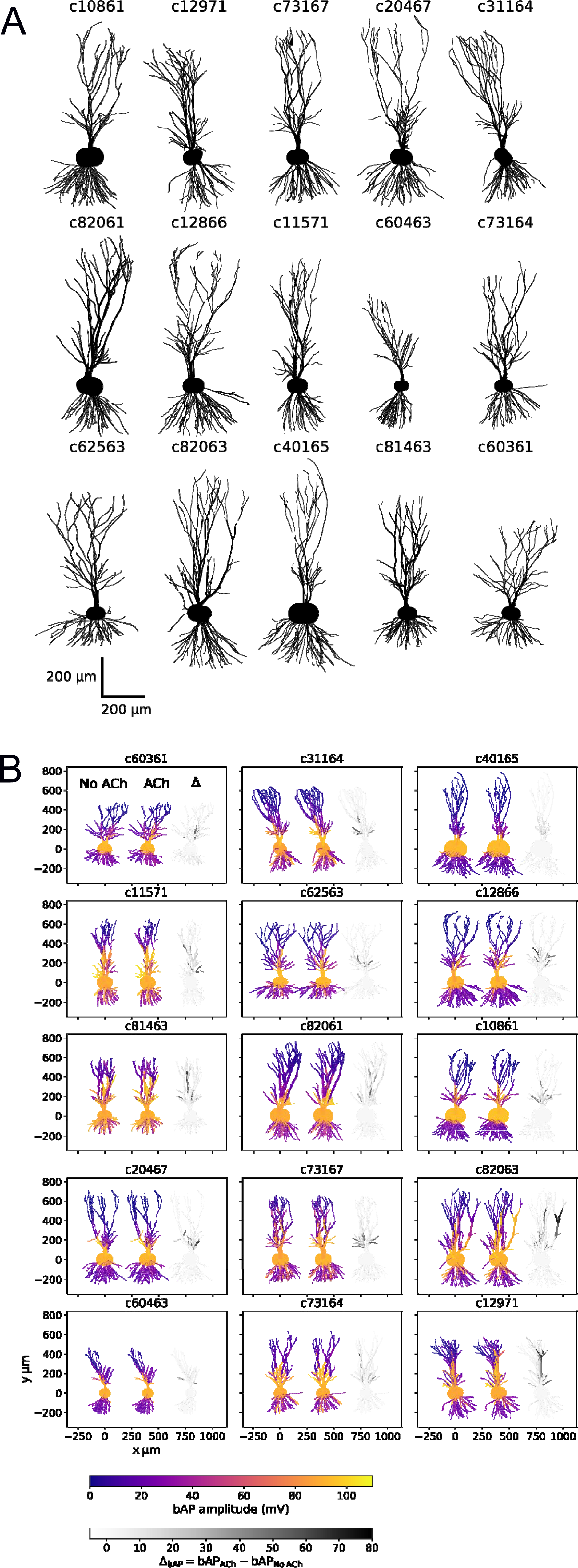
CA3 pyramidal cell morphologies used for biophysical modelling. A) All 15 cell morphologies plotted from NEURON spatial information. B) For each cell morphology, back-propagating action potential amplitude before (left) and after (middle) cholinergic modulation, and the difference in amplitude (right) are shown distributed across each CA3 pyramidal cell.

**Supplementary Figure 4:**
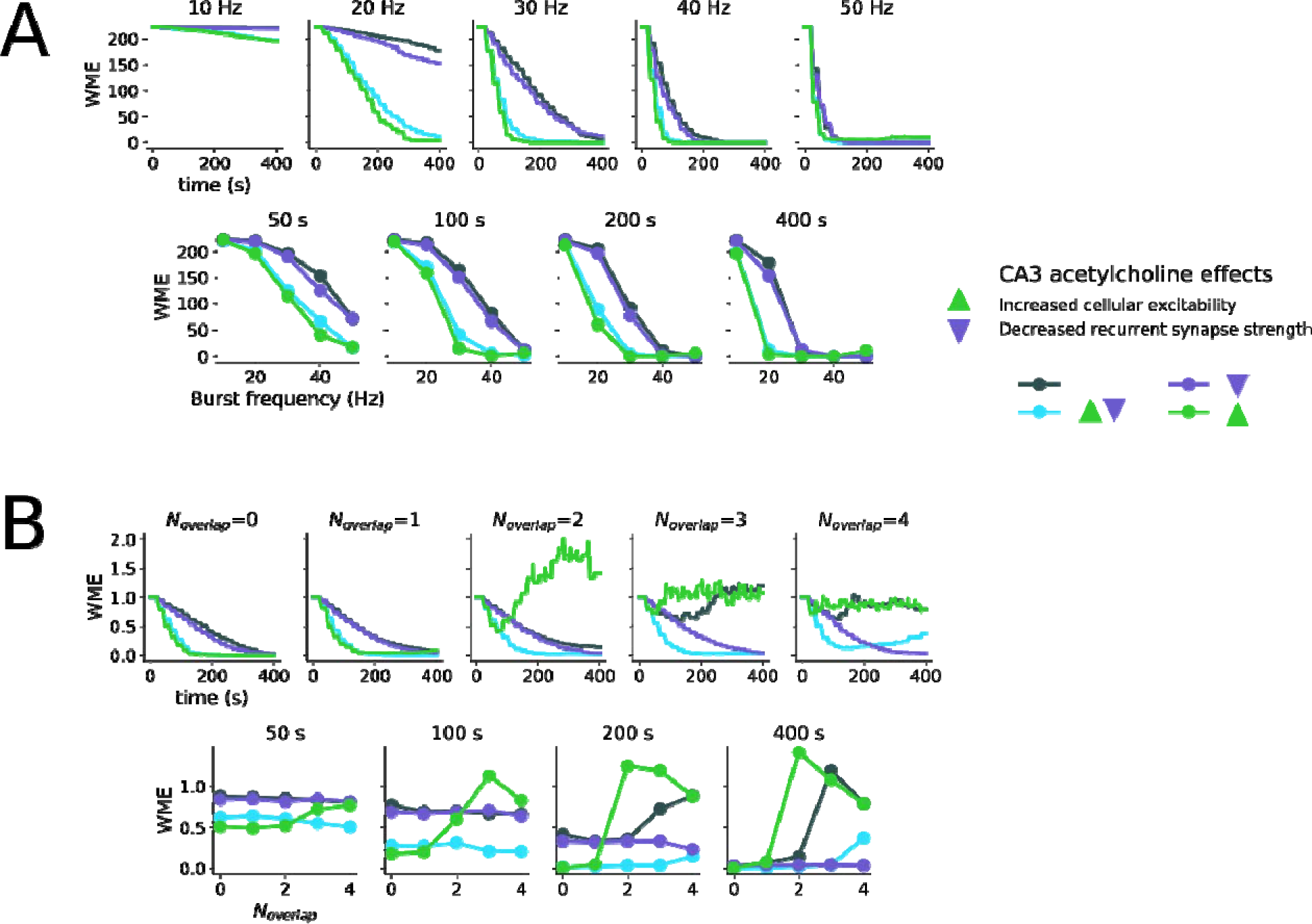
A) Slices of data shown in Figure 6C along frequency (top) and time (bottom) axes. B) Slices of data shown in Figure 7C along overlap (top) and time (bottom) axes. Colour coding for plots is indicated in the legend representing inclusion of different effects of acetylcholine in CA3.

**Supplementary Figure 5:**
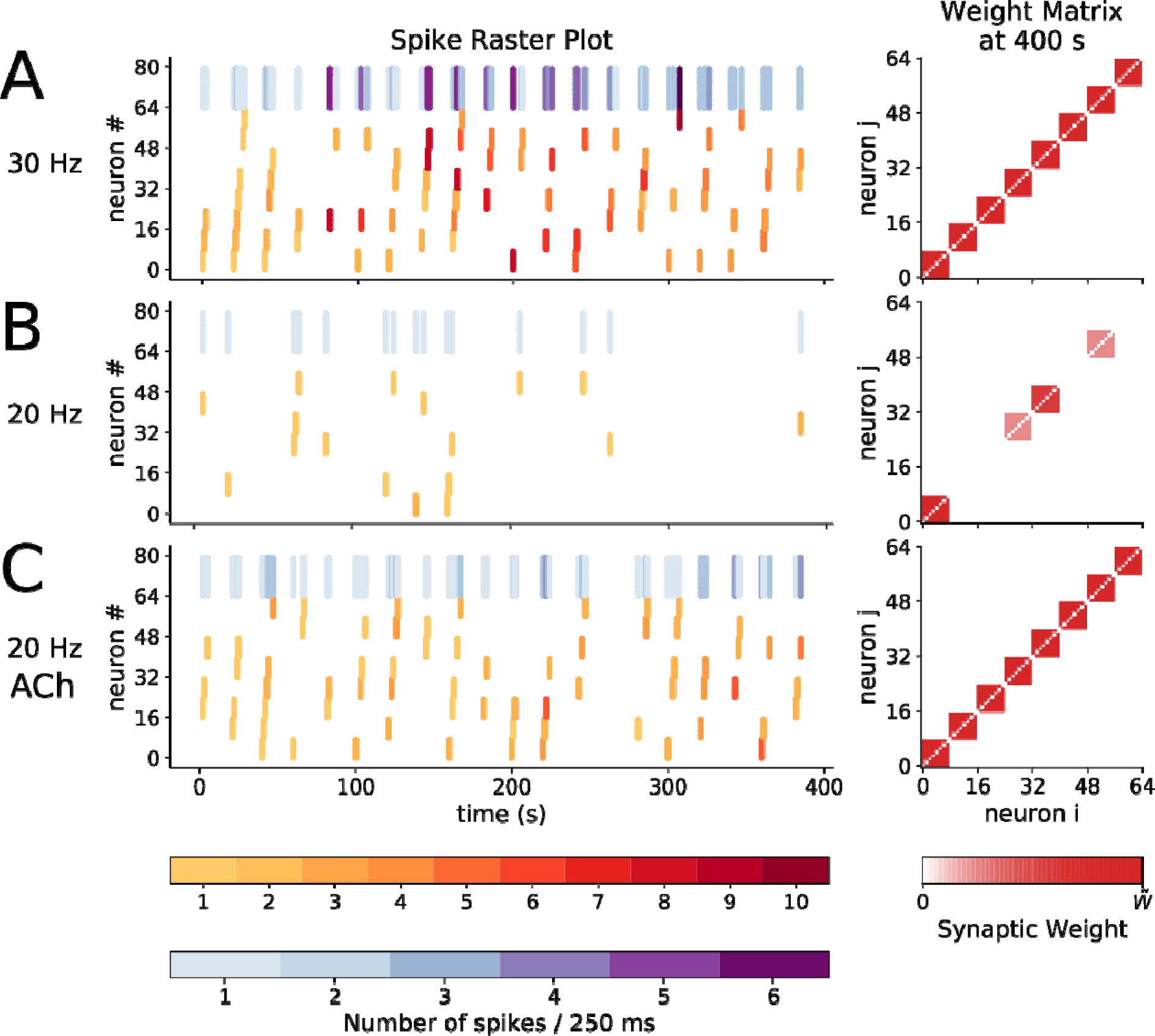
Model CA3 spiking activity (left) and resulting pyramidal cell weight matrix (right) with mossy fibre bursting at 30 Hz (A), 20 Hz (B), and 20 Hz with cholinergic modulation (C).

**Supplementary Figure 6:**
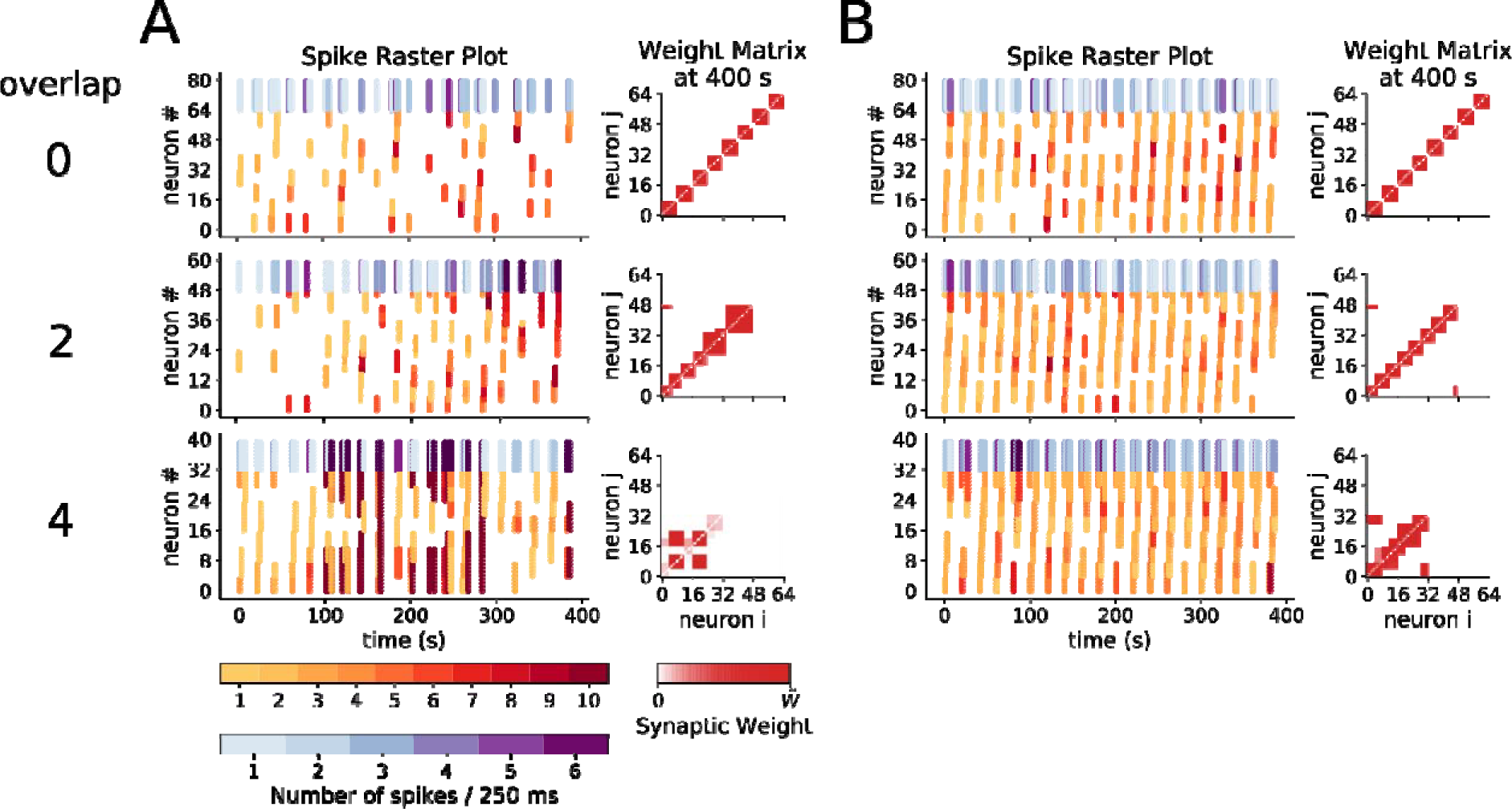
Model CA3 spiking activity and resulting weight matrix without cholinergic modulation (A) and with cholinergic modulation (B) with 0 (top), 2 (middle), and 4 (top) cells overlapping between ensembles.

